# Chromatin-associated MRN complex protects highly transcribing genes from genomic instability

**DOI:** 10.1101/2020.05.27.118638

**Authors:** Kader Salifou, Callum Burnard, Poornima Basavarajaiah, Marion Helsmoortel, Victor Mac, David Depierre, Giuseppa Grasso, Céline Franckhauser, Emmanuelle Beyne, Xavier Contreras, Jérôme Dejardin, Sylvie Rouquier, Olivier Cuvier, Rosemary Kiernan

**Author notes:** equal contribution (Kader Salifou, Callum Burnard, Poornima Basavarajaiah).

## Abstract

The MRN-MDC1 complex plays a central role in the DNA damage response (DDR) and repair. Using Proteomics of Isolated Chromatin Fragments (PICh), we identified DDR factors, such as MDC1, among those that become highly associated with a genomic locus upon transcriptional activation. Purification of endogenous MDC1, in the absence of exogenous DNA damage, revealed its interaction with factors involved in gene expression and co-transcriptional RNA processing, in addition to DDR factors. ChIP-seq analysis showed that MDC1 interacting factors, MRE11 and NBS1 subunits of MRN, were co-localized throughout the genome and notably at TSSs and gene bodies of actively transcribing genes. Blockade of transcriptional elongation showed that binding of MRN was dependent on the RNAPII transcriptional complex rather than transcription *per se*. Depletion of MRN increased RNAPII abundance at TSSs and gene bodies of MRE11/NBS1-bound genes. Prolonged exposure of cells to either MRE11- or NBS1-depletion led to single nucleotide polymorphism formation across actively transcribing, MRE11 or NBS1 target genes. These data support a model by which association of the MRN complex with the transcriptional machinery constitutively scans active genes for transcription-induced DNA damage to preserve the integrity of the coding genome.

## Introduction

Execution of the appropriate transcriptional programme is fundamental during growth, development and response to environmental stimuli. Although an essential DNA-dependent process, transcription comes at a cost. It can induce double strand breaks (DSBs) that are a deleterious form of DNA damage and can compromise the genome if erroneously repaired. DSBs are predominantly repaired by two canonical pathways: non-homologous end-joining (NHEJ) or homologous recombination (HR). HR involves extensive DNA end resection and uses an intact copy of the damaged locus to produce error-free repair. NHEJ ligates broken DSB ends with no or limited processing and frequently results in genomic mutations. Both pathways compete to repair a DSB (*1*) and mechanisms underlying pathway choice are not entirely clear. The nuclease activities of MRE11, a subunit of the MRE11-Rad50-NBS1 complex (MRN), are a critical determinant of pathway choice (*2*). MRE11 endonuclease activity initiates resection, thereby licensing HR. The exonuclease activities of MRE11 and EXO1/BLM bidirectionally resect toward and away from the DNA end, which commits to HR and disfavours NHEJ.

Recent studies have identified the local nuclear environment and epigenetic landscape as determinants of DNA repair pathway usage. DNA repair at actively transcribed regions occurs through the HR pathway whereas non-coding regions are more frequently repaired by NHEJ (*3-5*). Nuclear organization has emerged as a key parameter in the formation of chromosomal translocations, implying that DSBs cluster into higher order structures. Indeed, microscopy-based techniques and more recent chromosome conformation capture assays have shown that DSBs are mobile and form higher order structures detected as clusters (*6-9*). Initially thought to be factories for DNA repair, recent data have suggested that clustering may protect DSBs from spurious pairing to suppress translocations. Thus, DSBs in actively transcribed chromatin are not repaired in G1 but remain in clusters until HR can proceed in G2 (*7*). DSB clustering depends on several DNA repair factors including the MRN complex (*6, 7, 9*). Thus, a model emerges in which DSBs in active chromatin are sequestered in higher order structures that permit the HR repair pathway to proceed under conditions that minimize the formation of deleterious translocations. However, the molecular mechanism by which HR is chosen as a predominant pathway for DSB repair in transcriptionally active regions remains unclear. Furthermore, most studies have been performed by inducing DNA damage using irradiation, chemicals or artificially cutting the genome. Mechanisms involved in endogenous transcription-associated DNA repair remain poorly understood.

To address the recruitment of DNA damage repair (DDR) factors at transcriptionally active regions, we used an unbiased proteomic approach, Proteomics of Isolated Chromatin fragments (PICh) (*10*), of the inducibly transcribed HIV-1 promoter. Upon transcriptional activation, DNA repair factors were among the most recruited interactants. In particular, Mediator of DNA Damage Checkpoint (MDC1) that associates with the MRN complex was recruited specifically following activation of transcription. Identification of the interactome of endogenous MDC1 by tandem affinity purification followed by mass spectrometry similarly identified numerous factors implicated in transcription and co-transcriptional RNA processing. Co-immunoprecipitation analysis showed that MDC1 interacts with RNAPII and co-activators, such as P-TEFb. ChIP-seq analysis of MRN subunits, MRE11 and NBS1, revealed an association with genes, specifically those that were transcriptionally active. MRE11 and NBS1 demonstrated co-variation with RNAPII upon heat shock-induced transcriptional activation. Blockade of transcriptional elongation altered the profile of NBS1 to mirror that of RNAPII suggesting that MRN is recruited through its association with the transcriptional machinery, rather than the level of transcription *per se*. Following MRE11- or NBS1-depletion, the occurrence of single nucleotide polymorphisms (SNPs) was highly correlated to the binding of MRE11, NBS1 or RNAPII. These data suggest that the MRN complex, through its interaction with the transcription machinery, scans active genes for transcription-induced DNA damage, which may favour repair through the HR pathway at actively transcribed regions.

## Results

### DDR factors are Associated with Active Transcription

We investigated the recruitment of DDR factors during transcription using an unbiased proteomic method, Proteomics of Isolated Chromatin Fragments (PICh) (*10*) of the HIV-1 promoter region, which becomes highly transcribed upon addition of its cognate transactivator, Tat. PICh was performed using chromatin extracts of cells containing tandem integrated copies of the HIV-1 LTR linked to MS2 binding sites (*11*), in the presence or absence of the Transactivator protein, Tat (fig. S1A, B). Mass spectrometry identified 438 proteins associated with the locus in the control condition and 622 proteins associated with the locus upon transactivation by Tat. Upon transcriptional activation, 58 proteins were highly gained (FC >7), 294 were moderately gained (FC >2, <7), 29 were moderately lost (FC <-2, > −7) and 3 were highly lost (FC <-7) (Fig. 1A and table S1). Gene ontology revealed a number of pathways that were enriched in cells expressing Tat, compared to control untreated cells (fig. S1C). As expected, factors involved in gene expression were among the most highly gained. Interestingly, a number of factors involved in DNA repair were associated with HIV-1 chromatin specifically upon transactivation of transcription with Tat. Among these, Mediator of DNA damage Checkpoint (MDC1) was highly represented (Fig. 1A). To validate the interaction of DDR factors with actively transcribed HIV-1, we performed chromatin immunoprecipitation (ChIP) in cells containing an integrated HIV-1 LTR linked to a luciferase reporter, as described previously (*12*). Tat-induced activation of HIV-1 transcription increased recruitment of RNAPII as expected (fig. S1D). Similarly, association of DDR factors MDC1, MRE11, and Ku80 were also significantly enhanced in the presence of Tat (fig. S1D), in agreement with the recruitment of DDR factors upon transcriptional activation identified by PICh.

**Fig. 1.**
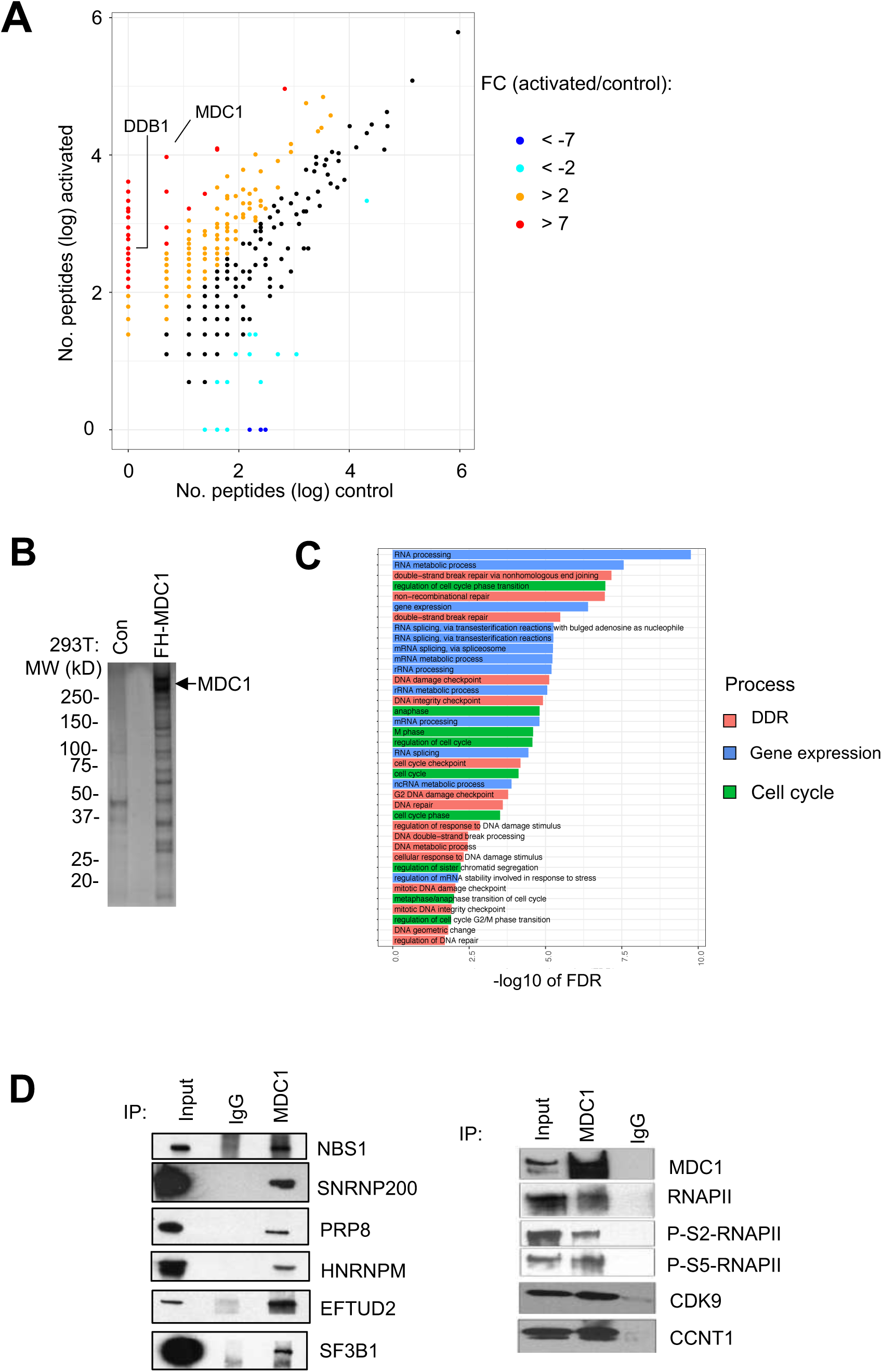
DNA repair factors are associated with active transcription. **(A)** Scatter plot representation of gain or loss of proteins associated with a tandem integrated HIV-1 minigene in the presence (activated) or absence (control) of Tat as indicated. **(B)** SDS-PAGE analysis followed by silver staining of eluates of tandem affinity purified nuclear extracts from HEK293T expressing Flag-HA fused to endogenous MDC1 (FH-MDC1) or control cells (con). **(C)** Gene Ontology analysis of FH-MDC1 interactome. **(D)** Co-immunoprecipitation analysis of MDC1 performed using the antibodies indicated on the figure.

These data indicated that DDR factors such as MDC1 and MRN may be closely linked with active transcription. To further explore this, we identified the interactome of MDC1 in the absence of exogenously induced DNA damage. Using Crispr-Cas9 genome editing technique, a Flag-HA tag was appended to the N-terminus of endogenous MDC1 protein in HEK-293T cells. Tandem affinity purification of Flag-HA MDC1 was performed, as described previously (*12-14*) (Fig. 1B). Gene ontology analysis of the interactants identified association of MDC1 with factors in the DNA damage repair pathway and the cell cycle, as expected (Fig. 1C). Interestingly, MDC1 interactants were also implicated in gene expression pathways (Fig. 1C, table S2). Association of endogenous MDC1 with RNA processing factors, such as SNRNP200, PRP8, HNRNPM, EFTUD2 and SF3B1 was validated by co-immunoprecipitation analysis (Fig. 1D, left panel). Interaction between MDC1 and RNAPII (total and CTD phosphorylated forms) as well as subunits of P-TEFb, CDK9 and cyclin T1, was also detected by co-immunoprecipitation analysis (Fig. 1D, right panel), confirming a previous report (*15*). Moreover, immunoprecipitates of cyclin T1 contained MDC1 and MRN subunits, MRE11 and NBS1 (Fig. S1E). Furthermore, MDC1 was specifically associated with the small, active P-TEFb complex, as shown by glycerol gradient analysis (fig. S1F). These data show the highly transcribing HIV-1 locus recruits DDR factors such as MDC1 and MRN and conversely, that DDR factor MDC1 is physically associated with factors implicated in transcription and RNA processing. Taken together, they reveal that the transcription and DDR pathways are intimately linked at the biochemical level through physical interactions.

### MRN Complex is Associated with Chromatin

Given its interaction with factors implicated in RNAPII transcription, we next sought to determine the profile of chromatin-associated MDC1/MRN complex across the coding genome. Anti-MDC1 antibodies proved unsuitable for ChIP-seq. We therefore performed ChIP-seq analysis of MRN subunits, NBS1 and MRE11, as well as RNAPII in cells in the absence of exogenous DNA damage. The average profile of signal for both MRE11 and NBS1 at active genes resembled that of RNAPII, although the amount of signal detected was lower (Fig. 2A). While the highest accumulation of MRE11 and NBS1 ChIP-seq reads occurred near the TSS, significant levels of binding were also detected in the gene body and downstream of the TES (Fig. 2A). The binding profile of MRE11 and NBS1 on representative genes, RBM17 and MED23, is shown in Fig. 2B. More than 50,000 peaks were detected throughout the genome for either MRE11 or NBS1, and 110,000 peaks detected for RNAPII. Although the highest signal for both MRE11 and NBS1 was found around TSSs (Fig. 2A, B), the number of peaks overlapping TSSs represented 2.19% and 1.44%, respectively (Fig. 2C). MRE11 and NBS1 were also detected on transcription end/termination sites (TESs; < 2 %) and enhancers (3.16 % and 1.74 %, respectively) (Fig. 2C). The majority of MRE11 and NBS1 peaks were found on gene bodies (approximately 52 %), with a distribution along gene bodies similar to that of RNAPII (fig. S2).

**Fig. 2.**
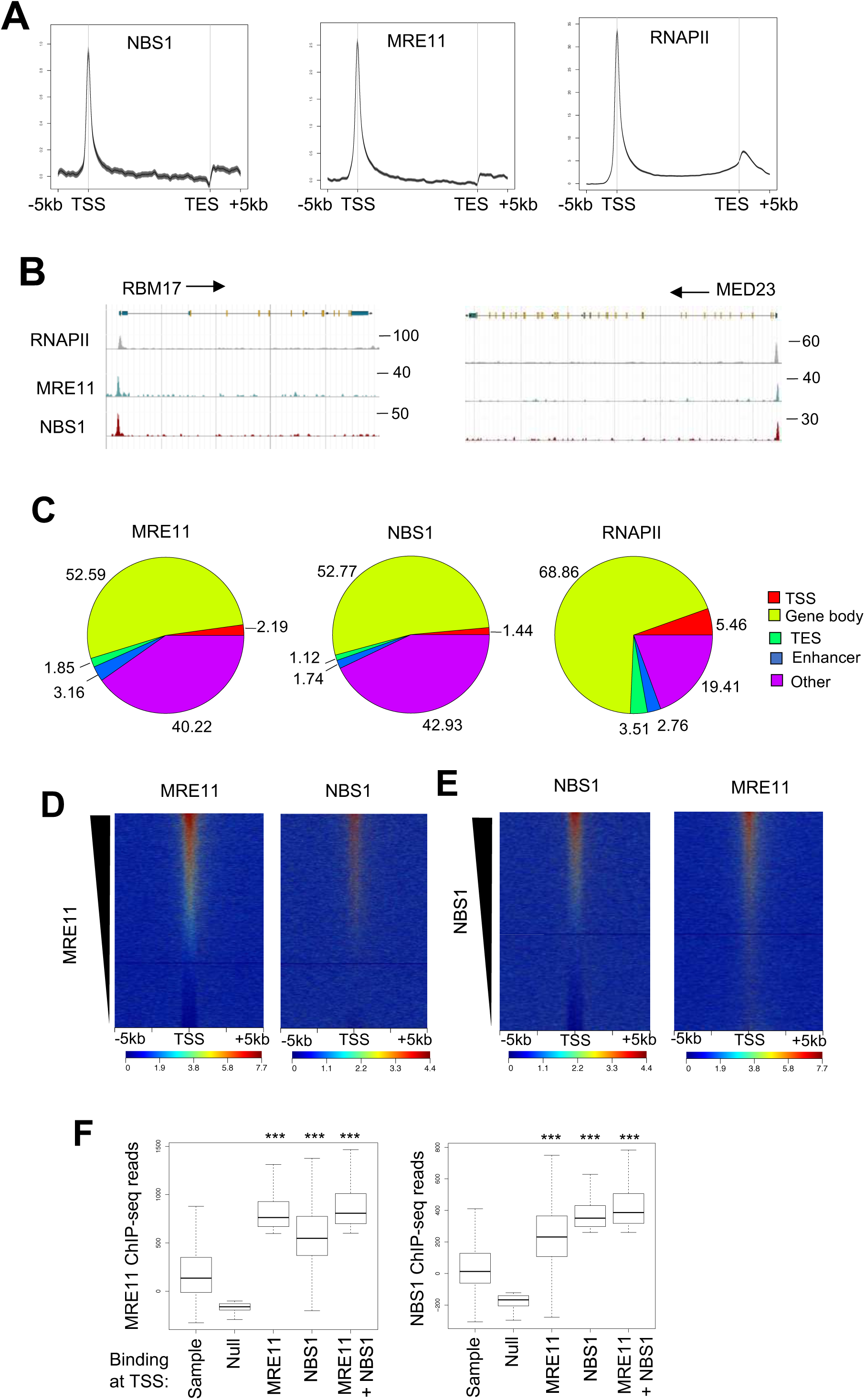
MRN Complex is Associated with Chromatin. **(A)** Average density profiles of MRE11, NBS1 and RNAPII ChIP-seq reads across genes, +/- 5 kB. **(B)** Chip-seq tracks of representative genes showing MRE11, NBS1 and RNAPII ChIP-seq signal as indicated. **(C)** Pie charts showing the genomic distribution of ChIP-seq peaks of MRE11, NBS1 and RNAPII, as indicated. **(D and E)** ChIP-seq heatmaps centered and rank-ordered on MRE11(D) or NBS1 (E). Chip-seq reads of NBS1 (D) or MRE11 (E) were plotted respecting the same ranking. **(F)** Box plots of ChIP-seq reads of MRE11 or NBS1 at genes highly bound at the TSS by MRE11, NBS1, both or neither, as indicated, compared to a random sample of genes (*** p < 0.001, Wilcoxon test), (n=708,479,1631,1631 and 708, respectively)

Further genomic analysis of the localization of MRN on chromatin was performed by ranking genes according to the binding of MRE11 at TSSs (number of ChIP-seq reads, highest to lowest), as shown in Fig. 2D, left panel. Ranking genes by MRE11 reads also efficiently sorted the NBS1 signal (Fig. 2D, right panel). Similarly, gene ranking based on NBS1 read intensity at the TSS also efficiently ranked the MRE11 signal (Fig. 2E), highlighting a good correlation in their loading onto genes. Next, genes were grouped based on high TSS binding of either MRE11 or NBS1, both factors together, or neither factor. As expected, genes highly bound by MRE11 had significantly more ChIP-seq reads for this factor compared to the random sample group (Fig. 2F, compare box 3 with 1; p < 1e-20). The group of genes that were highly bound by NBS1 also had a significantly high number of MRE11 reads at the TSS compared to the sample group (p <1e-20; left boxplot, compare box 4 with 1). Moreover, the highest number of number of MRE11 reads were found at co-bound genes when compared to the sample group (left boxplot, compare box 5 with 1; p <1e-20), or to TSSs highly bound by MRE11 only (p <1e-7; left boxplot, compare box 5 with 3). The same trend was observed for NBS1 binding (Fig. 2B, right boxplot). TSSs highly bound by NBS1 showed a high number of NBS1 reads as expected (compare box 4 with 1; p <1e-20), while TSSs highly bound by MRE11 also showed significantly high binding of NBS1 compared to the random sample group of genes (compare box 3 with 1; p <1e-20). As observed for MRE11, the highest number of NBS1 reads were found at genes that were co-bound by MRE11 (compare box 5 to 4; p <1e-13). These data indicate that MRE11 and NBS1 co-associate with a subset of human genes, likely in the context of the MRN DNA repair complex. They furthermore support proteomic data shown in Fig. 1 that uncovered biochemical interactions between MDC1/MRN complex and factors implicated in transcription and co-transcriptional mRNA processing.

### MRN Complex is Associated with Actively Transcribed Regions

Since MRE11 and NBS1 were found to be highly associated with TSSs and gene bodies, we wished to further investigate their association with transcription. First, genes were ranked according to the number of RNAPII ChIP-seq reads surrounding the TSS (+/-1 kb), from highest to lowest, which likely reflects the transcriptional activity of each gene. Next, respecting the same ranking, ChIP-seq reads for MRE11 and NBS1 were plotted. The resulting heatmaps (Fig. 3A) show that ranking genes by RNAPII also efficiently ranked both MRE11 and NBS1. Similarly, the averaged density profiles showed that MRE11 and NBS1 were more associated with genes having high RNAPII at the TSS, than those having low RNAPII (Fig. 3A, top panels).

**Fig. 3.**
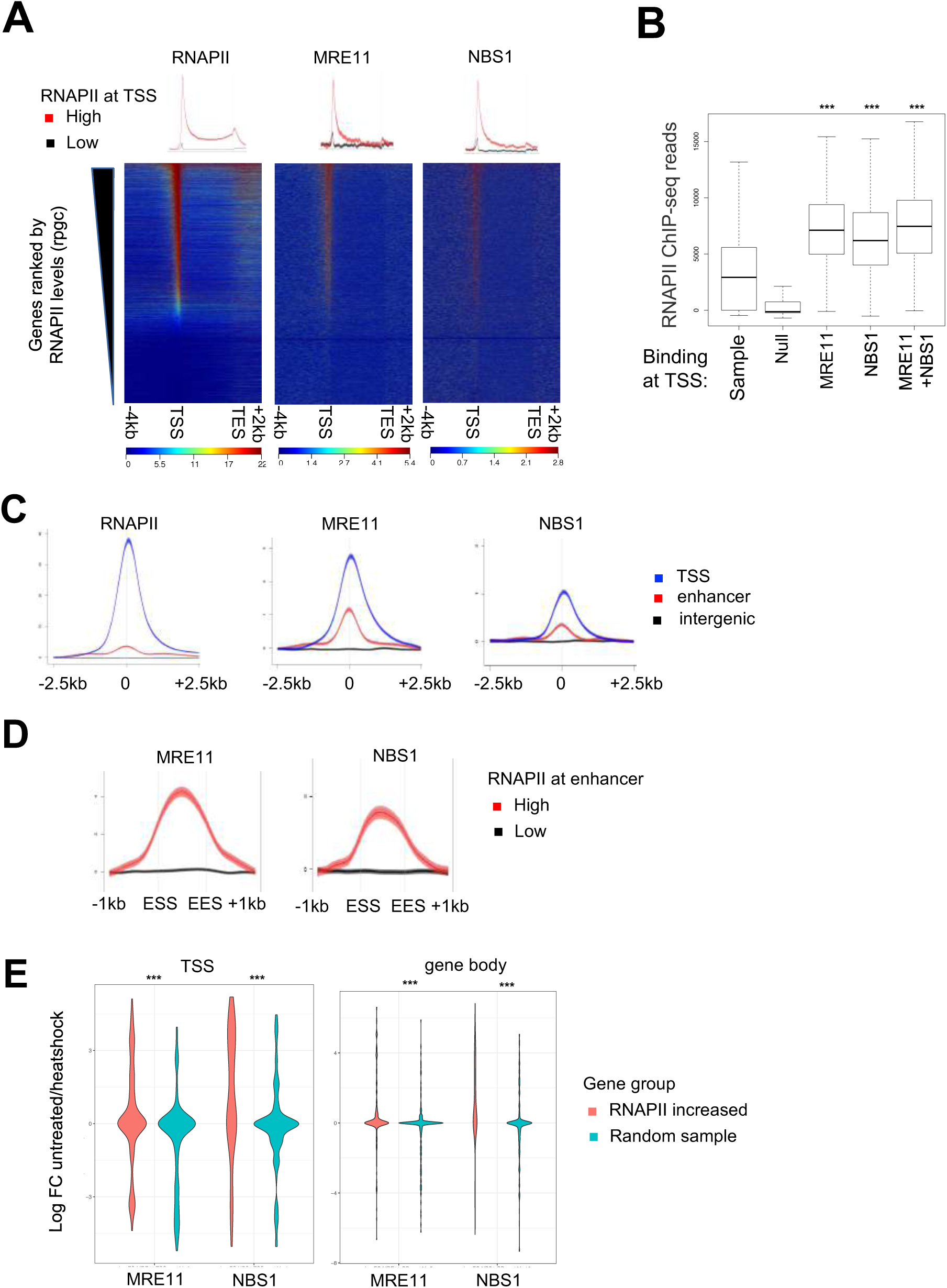
MRN Complex is Associated with Actively Transcribed Regions. **(A)** ChIP-seq heatmaps rank-ordered on RNAPII ChIP-seq reads at the TSS, and centered on genes, - 4 kB/+2 kB. Chip-seq reads of MRE11 or NBS1, as indicated, were plotted respecting the same ranking. Average density profiles of ChIP-seq reads of RNAPII, MRE11 or NBS1 at the highest ranked and lowest ranked RNAPII genes are shown above the heatmaps. **(B)** Box plots of RNAPII ChIP-seq reads at genes highly bound at the TSS by MRE11, NBS1, both or neither, as indicated, compared to a random sample of genes (*** p < 0.001, Wilcoxon test), (n=708,479,1631,1631 and 708, respectively). **(C)** Average density profiles of RNAPII, MRE11 or NBS1 ChIP-seq centered on TSS, enhancers or intergenic regions, +/- 2.5 kB. **(D)** Average density profiles of ChIP-seq reads of MRE11 or NBS1 at enhancers +/- 1 kB having the highest ranked and lowest ranked RNAPII ChIP-seq reads, as indicated (ESS, enhancer start site, EES, enhancer end site). **(E)** Violin plots showing changes (Log FC) in MRE11 or NBS1 ChIP-seq reads, as indicated, at the TSS and gene body of genes at which RNAPII ChIP-seq reads were increased in heat shocked samples compared to non-treated controls, compared to a random sample group of genes in the same conditions, as indicated (*** p < 0.001, Wilcoxon test).

We next analyzed the amount of RNAPII at the TSS of genes highly bound by MRE11, NBS1 both, or neither factor (null) compared to a random sample group of genes (sample). Genes highly bound by either NBS1 or MRE11, alone or together, had significantly higher levels of RNAPII at the TSS compared to the random sample group of genes (Fig. 3B; boxes 3, 4 and 5, compared to box 1; ps > 1e-20). The association of MRE11 and NBS1 at the TSS of genes having varying amounts of RNAPII at the TSS was validated by ChIP-qPCR. The abundance of MRE11 or NBS1 largely mirrored that of RNAPII (fig. S3A). Furthermore, intersection analyses of ChIP-seq data showed a preferential co-localization of MRE11/NBS1 ChIP-seq peaks with active genes as compared with inactive genes. Considering the peaks of each factor found at TSSs, 84% of MRE11 peaks (n=1597), 79% of NBS1 peaks (n=687) and 88% of MRE11+NBS1 peaks (n=672) were found at active TSSs. Since RNAPII is also found at active enhancers, we asked whether MRE11 and NBS1 might associate with these regions. Compared to TSSs and random intergenic regions, both MRE11 and NBS1 were localized to enhancers (Fig. 3C). Similar to TSSs, MRE11 and NBS1 were detected at enhancers with high levels of RNAPII, which are likely active enhancers, and were poorly associated with enhancers having low levels of RNAPII (Fig. 3D). Taken together, these data indicate the MRE11 and NBS1 subunits of MRN complex are associated with actively transcribing regions in a manner that appears to be proportional to that of RNAPII.

We next tested whether the MRN complex co-associates with RNAPII following induction of a specific transcriptional programme. To this end, cells were heat-shocked to induce the expression of a subset of genes. ChIP-seq of RNAPII, MRE11 and NBS1 was performed in heat-shocked and untreated cells. MRE11 and NBS1 binding was analyzed at genes induced upon heat shock, that is, those showing an increase in RNAPII association. As shown in Fig. 3E, both MRE11 and NBS1 became significantly associated with the TSS (left panel) and gene body (right panel) of upregulated genes compared to a sample group of genes. On the other hand, alpha-amanitin treatment was used to diminish RNAPII association with genes. As shown in fig. S3B, alpha-amanitin treatment induced loss of RNAPII as well as both MRE11 and NBS1 at target genes. Taken together, these data confirm that the binding of MRN tightly correlates with that of actively transcribing RNAPII.

### MRN Association with Chromatin Depends on RNAPII rather than Transcription Levels

Association of the MRN complex with actively transcribing regions is highly correlated to that of RNAPII (Fig. 3). However, it is not clear whether MRN is recruited through its interaction with the transcriptional machinery (Fig. 1) (*15*) or during the process of transcription. To distinguish between these possibilities, we sought to analyze MRN association under conditions where the abundance of RNAPII is not strictly correlated with the amount of transcription. To do so, RNAPII transcriptional elongation was blocked using the adenosine analogue 5,6-dichloro-1-β-D-ribofuranosylbenzimidazole (DRB). While DRB abolishes transcription in the coding region, it does not greatly affect the amount of transcriptional initiation. Consequently, DRB treatment causes RNAPII to accumulate at the 5’ end of genes disproportionately to the amount of transcription at the site. After 1 hour DRB treatment, RNAPII accumulated near the TSS and diminished in the coding region and particularly at the 3’ end of genes in the termination window, indicaing that RNAPII elongation was efficiently inhibited (Fig. 4A). A longer blockade with DRB (3-hour treatment) caused RNAPII to accumulate at TSS boundaries, causing apparent accumulation near the start of the gene body and also at upstream regions. By 1 hour post-release, the profile of RNAPII had returned to that in untreated cells at the TSS and in the gene body, with a small reduction still evident at the TES.

**Fig. 4.**
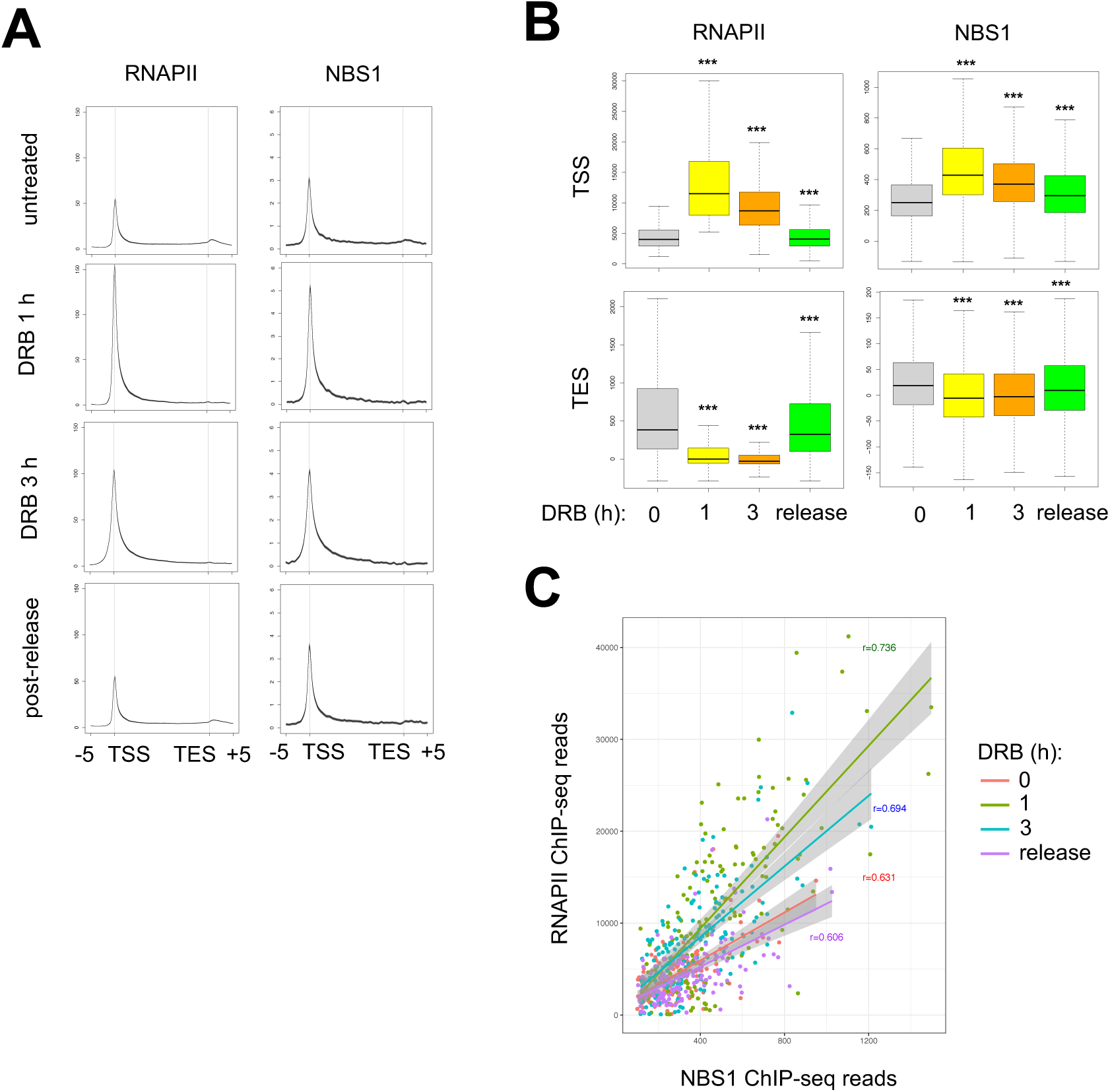
MRN association with chromatin is correlated with the amount of RNAPII rather than transcription. **(A)** Average density profiles of ChIP-seq reads of RNAPII and NBS1 across genes showing the highest increase in RNAPII reads at 1 h DRB treatment compared to a mock-treated control. ChIP-seq reads for RNAPII and NBS1 are shown at the same genes +/- 5 kB following treatment with DRB for 1h, 3h or 1 h post-release, as indicated. **(B)** Box plots of RNAPII and NBS1 ChIP-seq reads at the TSS and TES of regions shown in (A) in DRB-treated samples compared to nontreated controls (0) (***, p < 0.001, Wilcoxon test) (n=4924). **(C)** Scatter plot showing ChIP-seq reads of RNAPII and NBS1 at genes shown in (A) in untreated samples or samples treated with DRB, as indicated. The co-efficient of correlation (r) calculated for each condition is indicated on the graph.

Analysis of the profile of NBS1 under conditions of DRB treatment and release showed that it largely mirrored that of RNAPII, although the effects were more modest. The profiles showed the same qualitative effects, such as loss of signal at the TES, and leaking into TSS adjacent regions observed after 3h treatment. The change in profiles of NBS1 and RNAPII were also quantitatively significant at both the TSS and TES regions (Fig. 4B). Finally, we analysed the correlation between RNAPII and NBS1 association at individual genes for each condition. Association of the 2 factors with genes was positively correlated at each condition, with the highest correlation at 1 and 3 h DRB treatment (Fig. 4C). Thus, overall, although the effects measured for NBS1 are more modest than those for RNAPII, the similarities in their binding profiles and dynamics suggests that MRN is likely recruited through its association with the RNAPII transcription complex, rather than the amount of transcription *per se*.

### Binding of MRN Impacts the Transcriptional Output of Target Genes

We next wondered whether the presence of MRN impacts the level of transcription at its target genes. To test this, MRE11 and NBS1 were depleted by RNAi. Consistent with previous findings, knock-down of MRE11 induced loss of MRE11 and also destabilized NBS1 in cell extracts (fig. S4A), since MRE11 is required for stability of the MRN complex (*16*). In contrast, RNAi-mediated depletion of NBS1 did not have a significant impact on MRE11 (fig. S4A), as reported previously (*16*). Next, RNAi-depleted cells were analyzed by ChIP-seq for the recruitment of each factor and RNAPII. As expected, depletion of MRE11 led to a significant loss of MRE11 signal at genes highly bound at the TSS by MRE11, NBS1 or MRE11+NBS1 (fig. S4B, left panel, compare blue boxes to grey boxes). Consistent with the reduction of NBS1 in extracts following knock-down of MRE11, NBS1 binding to chromatin was also significantly reduced under the same conditions (fig. S4B, right panel, compare blue boxes to grey boxes). Similarly, depletion of NBS1 reduced NBS1 binding at TSSs highly bound by NBS1, alone or together with MRE11, (right panel, compare red boxes to grey boxes). Depletion of NBS1 also diminished MRE11 signal at genes highly bound by MRE11, alone or together with NBS1 (left panel, red boxes 9 and 15), supporting the idea of cooperative binding to chromatin.

We next analysed RNAPII association with genes in MRN-depleted and control cells. Interestingly, RNAPII accumulated over genes highly bound at the TSS by MRE11, NBS1 or both, following depletion of either MRE11 or NBS1 compared to a control knock-down (Fig. 5A). RNAPII binding was significantly increased at the TSS of genes highly bound by MRE11, NBS1 or both following loss of MRE11 or NBS1 (Fig. 5B, left and right panels, respectively, and fig. S5A) and across gene bodies (fig. S5B). Interestingly, depletion of either NBS1 or MRE11 altered RNAPII association at a common set of genes, not only at genes where RNAPII levels were increased (fig. S5C, right matrix, compare deciles 1 on X and Y axes), but also at genes where RNAPII association was decreased (fig. S5C, right matrix, compare deciles 10 on X and Y axes, p < 1e-20 by hypergeometric test). We next intersected the changes in RNAPII association with occupancy of MRE11 or NBS1. As shown in fig. 5C, genes showing the highest increase in RNAPII ChIP-seq reads (deciles 1 on X and Y axes) were also highly associated with MRE11 (left panel) or NBS1 (right panel). Although depletion of NBS1 or MRE11 led to loss of RNAPII at a common set of genes (deciles 10 on X and Y axes), these, in contrast, were not significantly bound by either factor.

**Fig. 5.**
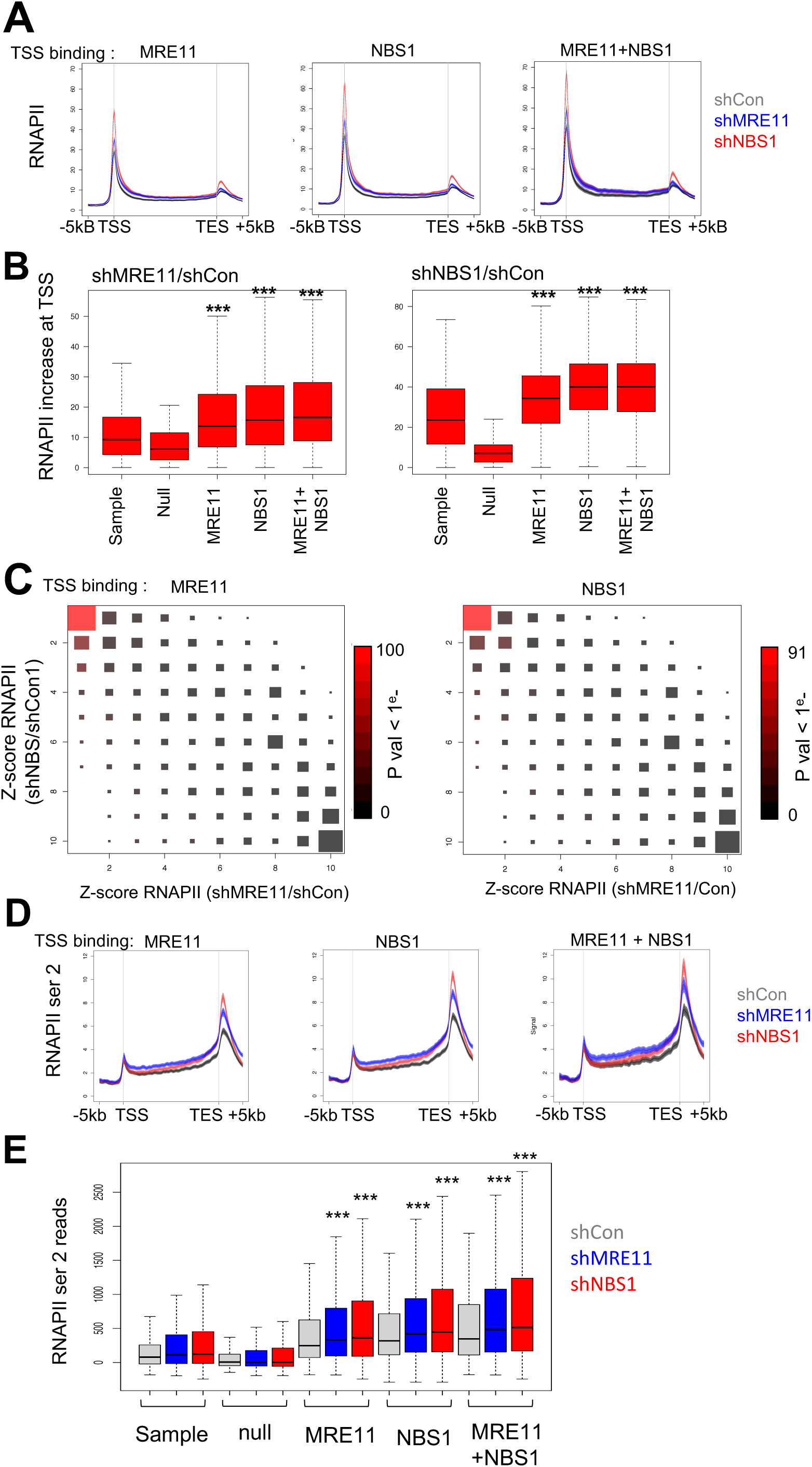
Binding of MRN Impacts the Transcriptional Output of Target Genes. **(A)** Average density profiles of RNAPII ChIP-seq reads across genes highly bound at the TSS by MRE11, NBS1 or both in samples treated with shRNA targeting MRE11, NBS1 or a non-targeting control, as indicated. **(B)** Box plots of RNAPII increase as z-scores at the TSS of genes highly bound at the TSS by MRE11, NBS1, both or neither, compared to a sample group of genes, following knock-down of MRE11 or NBS1, relative to a control knock-down. (***, p < 0.001, Wilcoxon test) (left box plot: n=443, 214, 1172, 1221 and 491, respectively, and p < 1e-12, p < 1e-20, p < 1e-18, respectively; right box plot: n=529, 176, 1519, 1548 and 625, respectively, and p < 1e-28, p < 1e-61, p < 1e-42, respectively) **(C)** Hinton diagram showing changes in RNAPII ChIP-seq reads at genes following knock-down of MRE11 or NBS1, relative to a control knock-down, ranked by deciles from the most up-regulated (1) to the most down-regulated (10), at genes highly bound at the TSS by either MRE11 (left diagram) or NBS1 (right diagram). The scale represents P-value for the intersection of RNAPII increase with TSS binding of MRE11 or NBS1 (10^−0^ to 10^−100^ and 10^−0^ to 10^−91^, respectively, Hypergeometric test). The size of the box represents the number of genes, as shown in Fig. S5C. **(D)** Average density profiles of phospho-Ser2 RNAPII ChIP-seq reads across genes +/- 5 kB highly bound at the TSS by MRE11, NBS1 or both in samples treated with shRNA targeting MRE11, NBS1 or a non-targeting control, as indicated. **(E)** Box plots of ChIP-seq reads of phospho-Ser2 RNAPII at the TES of genes highly bound at the TSS by MRE11, NBS1, both or neither, compared to a sample group of genes, following knock-down of MRE11 or NBS1 or a control knock-down, as indicated (***, p < 0.001, Wilcoxon test) (n=665, 398, 1631, 1631 and 665, respectively).

To determine whether the increase in RNAPII association with genes that occurred upon depletion of MRN reflected an increase in processive transcription, we determined the association of the elongating, serine 2 phosphorylated form of RNAPII (ser2) at genes highly bound by MRE1 and/or NBS1. Ser2 RNAPII accumulated over genes bound by MRE11, NBS1 or both, following depletion of either MRE11 or NBS1 compared to controls (Fig. 5D) in a significant manner (Fig. 5E). MRN was also associated with active enhancers (Fig. 3C, D). Analysis of RNAPII ChIP-seq showed that, like at genes, loss of MRE11 or NBS1 led to an increase in RNAPII association at the same regions (fig. S5D). However, in contrast to genes, Ser2 RNAPII ChIP-seq reads were not significantly increased upon depletion of the factors (P-val < 0.05). Therefore, MRE11 and NBS1 have a global impact on RNAPII levels genome-wide, frequently at the same regions. Upregulation of RNAPII occurred at MRN-bound regions, whereas down-regulation of RNAPII may be independent of MRN association. These data suggest that MRN modulates the transcriptional output of target genes.

### MRN Protects Actively Transcribed Regions from Genomic Instability

Given the key role of MRN in genomic stability, we next assessed single nucleotide polymorphisms (SNPs) by whole genome sequencing in cells exposed to depletion of MRE11 or NBS1 or a control depletion. We discarded common SNPs found in all samples, since these are likely to be false-positives due to the background genomic environment. Approximately 30,000 SNPs unique to shCon cells were identified while nearly 100,000 SNPs were detected in MRE11- or NBS1-depleted cells (Fig. 6A). Within genes, the majority of SNPs localized to gene bodies (Fig. 6A), which may reflect the genomic distributions of MRE11 and NBS1 (Fig. 2C). A ranking test by Gene Set Enrichment Analysis (GSEA) showed that SNPs found in MRE11- or NBS1-depleted cells could be readily predicted by the binding of MRE11 or NBS1 at TSSs (Fig. 6B). Furthermore, in keeping with the strict association between RNAPII and MRN occupancy at genes, SNP frequency was also well predicted by binding of RNAPII at the TSS. Therefore, binding of MRN over gene bodies preserves genes from mutations detected as SNPs. This highlights the key function of the MRN complex in maintaining genomic stability at actively transcribed regions.

**Fig. 6.**
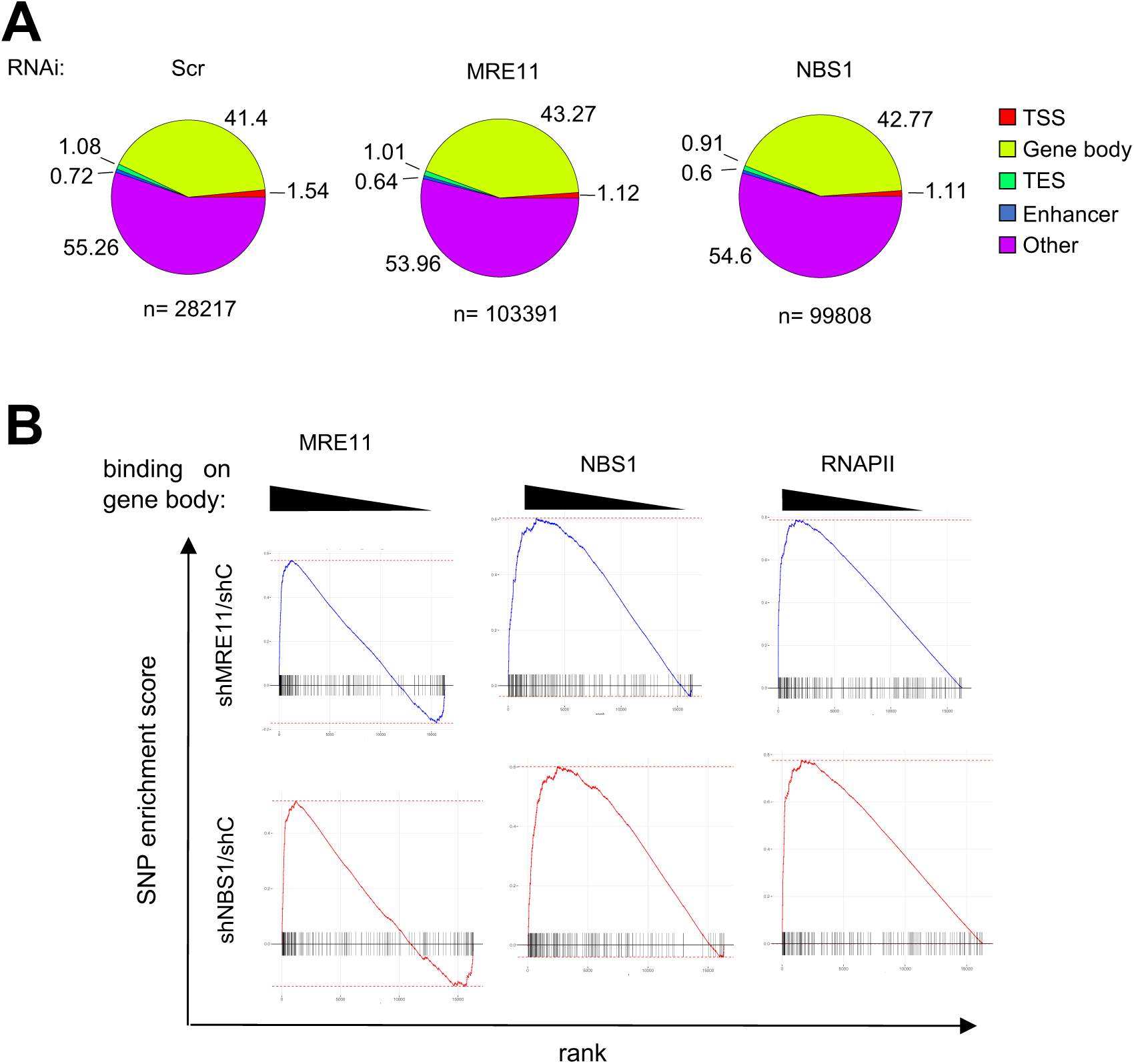
MRN Protects Actively Transcribed Regions from Genomic Instability. **(A)** Pie charts showing the genomic distribution of SNPs detected in samples following depletion of MRE11, NBS1 or a control, as indicated. **(B)** GSEA of SNPs detected in samples following depletion of MRE11, NBS1 compared to shCon sample, as indicated, ranked by ChIP-seq reads of MRE11, NBS1 or RNAPII in gene bodies, as indicated.

## Discussion

Recent data strongly point to a central role for the MRN complex in the resolution of transcription-associated DNA repair. However, mechanistic details about the recruitment of MRN to sites of transcription-associated DNA damage are not clear. Using the unbiased proteomic technique, PICh, we found that DNA repair factors were among the most recruited proteins that specifically interacted with the chromatin region during transcription of the inducible HIV-1 LTR. In particular, MDC1, which interacts with MRN complex, became highly associated with the actively transcribed locus and was largely absent prior to transcriptional stimulation. Furthermore, we identified the interactome of MDC1 in cells in the absence of exogenous DNA damage. Consistent with results obtained by PICh, MDC1 interacted with many factors implicated in transcription and co-transcriptional RNA processing, indicating an association between MDC1 and active transcription. Indeed, a robust interaction between MDC1 and RNAPII, as well as P-TEFb, was detected in the absence of exogenous DNA damage, suggesting a constitutive association with transcription. ChIP-seq revealed widespread localization of endogenous MRE11 and NBS1 across genes. Association of both MRE11 and NBS1 with genes was strongly correlated with transcriptional activity. Our data further suggest that MRN association with active genes is dependent on the presence of the RNAPII transcriptional complex, rather than the level of transcription *per se*.

While it is unclear precisely how MRN associates with the RNAPII complex, it is interesting to note that MDC1 interacts with topoisomerase II (TOP2) through its C-terminal BRCT domains (*16*). TOP2 is a component of the RNAPII transcriptional complex and is required for transcription through chromatin (*17*). Whether MDC1 associates with the RNAPII complex through TOP2 or another transcription-associated factor, MRN is found at the most highly transcribed genes, and its association with genes is highly correlated to that of RNAPII. Notably, upon depletion of either MRE11 or NBS1, MRN target genes accumulated DNA damage in an MRN and RNAPII-dependent manner. Taken together, these data suggest that MRN is associated with the RNAPII transcription complex probably to scan for transcription-associated DNA damage and initiate repair.

NHEJ and HR pathways compete for the repair of DSBs (*1*). The degree of chromatin compaction is thought to influence repair pathway choice. DSBs occurring in open chromatin undergo end resection and are predominantly repaired by HR, which results in faithful repair and suppresses dangerous mutations from arising in coding regions of the genome (*5, 7, 18*). The preference for employing HR to repair DSBs in transcribed regions is not due to cell cycle-dependent availability of factors. Increasing evidence indicates that DNA damage occurring on active genes is repaired through a specialized mechanism involving HR. The current model suggests that upon DNA damage the damaged site is targeted by MRN complex and undergoes the initial steps in HR, which disfavors repair of the site by NHEJ. Such MRN-marked sites subsequently pair and cluster into higher order structures to be repaired flawlessly during G2 phase of the cell cycle by the HR pathway. This mechanism preserves the fidelity of the genome in coding regions. How does the HR pathway outcompete NHEJ, which is constitutively available and actively repairs intergenic regions, at transcribed regions? We speculate that the physical link between MRN, which is required for commitment to HR, and RNAPII transcription complex, could in part explain how HR is selected over NHEJ at transcribed regions. Being a component of the transcription complex, MRN may have immediate access to transcription-induced DSBs, facilitating end resection and commitment to HR.

Several lines of evidence indicate that the presence of DSBs in coding regions leads to transcriptional arrest. For example, transcriptional silencing occurs on chromatin in the vicinity of DSBs in an ATM-dependent manner (*19*). Paradoxically, we observed that transcription was modestly increased in cells depleted of MRE11 or NBS1. There are several possible explanations for this apparently conflicting result. It has recently been shown that in response to UV irradiation, RNAPII is released from promoter-proximal regions into gene bodies to promote detection of DNA damage within genes (*20*). Thus, the increased RNAPII occupancy observed upon loss of MRN could also be a scanning mechanism to rapidly detect DNA damage within transcribed genes. Alternatively, the measured increase in transcription could be due to failure to establish HR at damaged sites in the absence of MRN. The use of HR to repair damage at active genes privileges genome fidelity at a slight cost to gene expression. Transcription of genes that have sustained DNA damage will be arrested until repair is completed in G2. Therefore, at the global level, expression of the gene will be somewhat diminished, but DNA repair will be flawless. In cells depleted of MRN, which is required to commit to HR, DNA repair most likely occurs by NHEJ. In this case, the cost to transcription will be minimal, as NHEJ operates throughout the cell cycle. However, repair will occur at the expense of genome fidelity, as NHEJ repair is characterized by the appearance of SNPs. In fact, this is the outcome in cells depleted of MRN. RNAPII occupancy at both the TSS and gene body was higher than in control cells. However, whole genome sequencing revealed that DNA repair was highly error-prone with the appearance of thousands of SNPs in the gene bodies of target genes. Furthermore, the number of SNPs detected was highly correlated to the abundance of MRN at the gene under control conditions. This scenario implies that MRN does not directly influence transcription, but that the absence of MRN predisposes to an alternative DNA repair pathway that is more favourable for gene expression, but at the cost of genome fidelity.

Finally, the detection of approximately 50,000 SNPs in gene bodies following prolonged exposure to MRN depletion, yet in the absence of any exogenous DNA damage, highlights the threat to genome fidelity posed by highly active transcription and the importance of MRN in suppressing genome mutation. The sources of transcription-associated DNA damage are likely numerous, ranging from increased accessibility during transcription, the generation of transcription-dependent RNA-DNA hybrids such as R loops, and the activity of DNA topoisomerases that relieve torsional stress by DNA strand passage reactions. Indeed, TOPO2 enzymatic activity generates a transient DSB. During such transient DSB formation, TOPO2 becomes covalently bound to the 5′ DNA end of the break, forming Top2-DNA cleavage complex intermediates (Top2cc). Failure to complete the religation step causes DSBs with associated Top2cc lesions. It was recently reported that failure to complete religation occurs much more frequently than previously thought, and that MRE11 nuclease activity plays a critical role in resolving such failed catalysis reactions (*21*). Thus, Top2cc lesions might be a potential source of the large number of SNPs found to be accumulated in MRN-deficient cells in this study.

## Materials and Methods

### Cell Culture, Treatment and Lentiviral Infection

HeLa and HEK-293T cells were grown in Dulbecco modified Eagle’s minimal essential medium (DMEM) (sigma-Aldrich D6429), supplemented with 10% FCS, and containing 1% Penicillin streptomycin (Sigma P4333). All cells were grown in a humidified incubator at 37 °C with 5% CO2. Where indicated, cells were subjected to heat-shock at 42°C for 1 hour, or treated with alpha-amanitin (Sigma-Aldrich, A2263) for 16 hours at 1 μg/ml. Cells were treated with 150 μM 5,6-dichloro-1-β-D-ribofuranosylbenzimidazole (DRB, Sigma-Aldrich D1916) or mock-treated with DMSO where indicated for 0, 1 or 3 h. After 3 h treatment, cells were washed, and incubated for a further 1 hour in fresh cell culture medium.

Production of shRNA-expressing lentiviral particles was performed as described previously (*12*) using plasmids expressing shRNAs targeting MRE11 (Sigma mission shRNA TRCN000338391), NBS1 (Sigma mission shRNA TRCN0000288622) or a non-targeting control (obtained through (Addgene, plasmid 1864), as shown in Table S3. For knockdown experiments, HeLa cells were transduced with lentiviral particles and harvested 4 days later, as described previously (*12*).

### Antibodies

Antibodies used in this study are shown in Table S4.

### CRISPR-Cas9 Mediated Editing of endogenous MDC1 gene

An sgRNA targeting the MDC1 gene around the ATG translation start site was cloned in pSpCas9 (BB)-2A-GFP plasmid (Addgene #48138). The plasmid was then transfected into HEK-293T cells along with a single stranded oligodeoxynucleotide (ssODN) (Table S5) harbouring the Flag-HA sequence flanked by homology sequences to MDC1 around the cleavage site. Single cells were isolated and amplified. HEK293T clones expressing Flag-HA MDC1 were identified by PCR and confirmed by Western blot using anti-HA and anti-Flag antibodies.

### Co-immunoprecipitation analysis

Co-immunoprecipitation was performed as described previously (*12*) using antibodies shown in Table S4.

### PiCh

Proteomics of Isolated Chromatin segments was performed as described previously (*10*) using U20S cells expressing HIV-LTR-MS2 (*11*). Sequences of probes used are shown in Table S5. Silver-staining was performed according to the manufacturer’s instruction (Silverquest, Invitrogen). Mass spectrometry was performed at Taplin facility, Harvard University, Boston, MA.

### MDC1 Protein Complex Purification

MDC1 complexes were purified from nuclear extracts (<u>10.17504/protocols.io.kh2ct8e</u>) of HEK-293T cells stably expressing Flag-HA-MDC1 by two-step affinity chromatography (dx.doi.org/10.17504/protocols.io.kgrctv6). Sequential Flag and HA immunoprecipitations were performed on equal amounts of proteins. Silver-staining was performed according to the manufacturer’s instructions (Silverquest, Invitrogen). Mass spectrometry was performed at Taplin facility, Harvard University, Boston, MA.

### Glycerol gradient sedimentation analysis

Separation of active and inactive P-TEFb complexes was performed as described previously (*22*). Briefly, glycerol gradients (10%–30%) were established by pipetting 2 ml of each of the glycerol fractions (10, 15, 20, 25, and 30% v/v) in buffer A (20 mM HEPES [pH 7.9], 0.3 M KCl, 0.2 mM EDTA, 0.1% NP-40) into centrifugation tubes (Beckman), 331372. Gradients were formed by standing for 6 h at 4 °C. Cells were lysed in 0.5 ml of buffer A (10 mM Tris-HCl [pH 7.4], 150 mM NaCl, 2 mM EDTA, 1% NP-40, 0.1% protease inhibitor) for 30 min at 4 °C. The lysates were centrifuged at 10,000*g* for 10 min and the supernatants were loaded into tubes with the preformed glycerol gradients. Protein complexes were then fractionated by centrifugation in an SW 41Ti rotor (Beckman) at 38,000 rpm. for 21 h. Fractions (0.5 ml) were collected, precipitated with trichloracetic acid and finally analyzed by immunoblotting with the appropriate antibodies.

### Chromatin Immunoprecipitation, library preparation and sequencing

Chromatin immunoprecipitation followed by high throughput sequencing (ChIP-seq) (*23*) was performed from HeLa cells using the ChIP-IT High Sensitivity® kit from Active motif (ref #53040) according to the manufacturer’s instructions. Each ChIP used 30 μg of chromatin along with 4 μg of antibody detecting MRE11, NBS1, RNAPII or phospo-Ser2 RNAPII (Table S4). ChIP-seq libraries were constructed using the Next Gen DNA Library Kit (Active Motif 53216 and 53264). Library quality was assessed using Agilent 2100 Bioanalyzer and Agilent High Sensitivity DNA assay. High throughput sequencing was performed by Sequence-By-Synthesis technique using a NextSeq 500 (Illumina) at Genom’ic facility, Institut Cochin, Paris.

### ChIP-qPCR

ChIP experiments were performed using the ideal ChIP-qPCR kit (Diagenode, ref C01010180) following the manufacturer’s instructions. Sequences of primers used for Real time quantitatitive PCR analysis are shown in Table S5.

### Whole Genome Sequencing

Genomic DNA was extracted from cells treated with lentiviral particles expressing shCon, shMRE11 or shNBS1 using the Qiagen DNeasy blood and tissue kit (ref 69504), according to the manufacturer’s instructions. To avoid RNA contamination, extracts were treated with RNAse A (Qiagen, 19101) according to the manufacturer’s instructions. Equal amounts of DNA were used for library preparation. Whole genome sequencing was performed by Novogene, Cambridge, UK.

### Bioinformatic Analysis

For PICh and mass spectrometry proteomics data analysis, uniquely mapping peptides were counted for each protein in each condition. Proteins whose abundance was greater than 7-fold that in the control condition were analysed using Gene Ontology enrichment analysis (*24, 25*), searching for enriched biological processes.

For analysis of ChIP-seq data, sequencing reads were first filtered, using fastq_illumina_filter, and quality control of filtered reads was performed using FastQC (http://www.bioinformatics.babraham.ac.uk/projects/fastqc/). Filtered reads were then aligned onto the HG38 genome (*26*) using the Burrows-Wheeler Aligner (BWA, http://bio-bwa.sourceforge.net/) with default parameters. The sorted BAM files generated by SAMtools (http://www.htslib.org), keeping only reads with a mapping quality at least 30, were then normalised by DeepTools’ (*27*) bamCoverage function, with a binsize of 10 bp. RPGC normalisation was applied, with an effective genome size of 2913022398, according to DeepTools’ user manual instructions. Files were then further normalised by subtracting an RPGC normalised input data file, using bigwigCompare. The input data used for normalisation was generated by averaging four inputs from separate assays, so as to minimise variability and biases that can be introduced during input normalisation.

From these normalised data files, peak calling was performed using NormR’s (https://doi.org/10.1101/082263) enrichR function, searching for enrichment of each BAM file of ChIP-seq reads against the input BAM file, using a False Discovery Rate (FDR) correction. Genomic Ranges (*28*) was then used to determine overlap between the peak range and genomic features of interest, such as genes with a transcription start site (TSS) and transcription termination site (TES) from GRCh38 and enhancers in HeLa S3 from ENCODE. Profile matrices were extracted from the normalised data files using DeepTools’ computeMatrix, using a bin size of 10 bp. For each gene, matrices for both TSS and TES were used, with 5 kb flanking each side of the feature, as well as gene bodies, which were scaled to 4 kb in length and with 4 kb flanking before the TSS and 2 kb flanking after the TES. For enhancers, the enhancer body was scaled to 500 bp, with 1kb flanking on either side. Using these profile matrices, quantification of normalised reads was calculated by summing the score of each appropriate bin for the feature. Unless indicated otherwise, this was from start to end for gene body and for enhancers, and 500 bp before and after TSS and TES.

RNAPII binding variation between conditions was calculated using z-scores, which were calculated as follows: 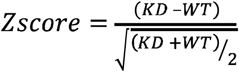 (i.e. difference weighted by mean signal), which transforms the distribution of variations into a normal distribution, allowing for better statistical interpretation of the variations.

Scatter plots, box plots, violin plots, pie charts and bar plots were created using either basic R plotting functions or ggplot2 (https://ggplot2.tidyverse.org) functions. Average binding profiles of proteins across genomic features of interest were generated using seqplots’ (DOI: <u>10.18129/B9.bioc.seqplots</u>) plotAverage function. Heatmaps on genomic features were created using genomation’s (DOI: <u>10.18129/B9.bioc.genomation</u>) gridHeat function, using the profile matrices generated by DeepTools. Gene set enrichment analysis was performed using fgsea (DOI: <u>10.18129/B9.bioc.fgsea</u>), and all enrichment plots were created using the plotEnrichment function of the package. Decile matrices were created using color2D matplot from the plotrix package (https://www.rdocumentation.org/packages/plotrix/versions/3.7-7). Genes ranked by z-score of RNAPII variation were split into deciles, for both shMRE11 vs shCon and shNBS1 vs shCon. Then, for each pair of deciles, the number of genes in the intersection of those two deciles was saved into a matrix, creating a 10×10 matrix of integers. The number of genes in each intersection was tested for significance by hypergeometric test, creating a 10×10 matrix of p-values. The genes at each intersection were also tested by hypergeometric test for significant enrichment in either top MRE11 or NBS1 binding levels (decile 1 of genes ranked by MRE11 or NBS1 binding).

For analysis of whole genome sequencing data, following processing by Annovar (*29*) on the sequencing platform, SNP location was then overlapped with gene bodies using Genomic Ranges functions, which yields a quantification per gene.

### Statistical Analysis

Data presented as histograms are shown as means ± SD. Comparison between two groups was analyzed by two-tailed Student’s *t* test, and asterisks represented significance defined as \**P* < 0.05, \*\**P* < 0.01, or \*\*\**P* < 0.001. Other statistical methods are described above in ‘bioinformatic analysis’.

## Acknowledgments

We thank Catherine Dargemont, and members of the Cuvier and Kiernan labs for helpful criticism, Juliette Hamroune and the Genom’ic platform for sequencing and Lisa Bluy for technical assistance.

## Author contributions

K.S, P.B, M.H. V.M. C.F. X.C. and S.R performed experiments and analysed data. C.B and D.D analysed data. R.K. and S.R. designed the experiments, O.C designed data analysis. R.K. wrote the manuscript with input from all authors.

## Data and materials availability

ChIP-seq data have been deposited at GEO (GSE 143591).

