## Supplementary material for "Chromatin-associated MRN complex protects highly transcribing genes from genomic instability": suppl info

**Supplemental Information**

**EXPERIMENTAL PROCEDURES**

**Cell Culture, Treatment and Lentiviral Infection**

HeLa and HEK-293T cells were grown in Dulbecco modified Eagle's minimal essential medium (DMEM) (sigma-Aldrich D6429), supplemented with 10% FCS, and containing 1% Penicillin streptomycin (Sigma P4333). All cells were grown in a humidified incubator at 37 °C with 5% CO2. Where indicated, cells were subjected to heat-shock at 42°C for 1 hour, or treated with alpha-amanitin (Sigma-Aldrich, A2263) for 16 hours at 1 μg/ml. Cells were treated with 150 μM 5,6-dichloro-1-β-D-ribofuranosylbenzimidazole (DRB, Sigma-Aldrich D1916) or mock-treated with DMSO where indicated for 0, 1 or 3 h. After 3 h treatment, cells were washed, and incubated for a further 1 hour in fresh cell culture medium.

Production of shRNA-expressing lentiviral particles was performed as described previously (Contreras et al., 2018) using plasmids expressing shRNAs targeting MRE11 (Sigma mission shRNA TRCN000338391), NBS1 (Sigma mission shRNA TRCN0000288622) or a non-targeting control (obtained through (Addgene, plasmid 1864), as shown in Table S3. For knockdown experiments, HeLa cells were transduced with lentiviral particles and harvested 4 days later, as described previously (Contreras et al., 2018).

**Antibodies**

Antibodies used in this study are shown in Table S4.

**CRISPR-Cas9 Mediated Editing of endogenous MDC1 gene**

An sgRNA targeting the MDC1 gene around the ATG translation start site was cloned in pSpCas9 (BB)-2A-GFP plasmid (Addgene #48138). The plasmid was then transfected into HEK-293T cells along with a single stranded oligodeoxynucleotide (ssODN) (Table S5) harbouring the Flag-HA sequence flanked by homology sequences to MDC1 around the cleavage site. Single cells were isolated and amplified. HEK293T clones expressing Flag-HA MDC1 were identified by PCR and confirmed by Western blot using anti-HA and anti-Flag antibodies.

**Co-immunoprecipitation analysis**

Co-immunoprecipitation was performed as described previously (Contreras et al., 2018) using antibodies shown in Table S4.

**PiCh**

Proteomics of Isolated Chromatin segments was performed as described previously (Dejardin and Kingston, 2009) using U20S cells expressing HIV-LTR-MS2 (Boireau et al., 2007). Sequences of probes used are shown in Table S5. Silver-staining was performed according to the manufacturer’s instruction (Silverquest, Invitrogen). Mass spectrometry was performed at Taplin facility, Harvard University, Boston, MA.

**Bioinformatic Analysis**

For PICh and mass spectrometry proteomics data analysis, uniquely mapping peptides were counted for each protein in each condition. Proteins whose abundance was greater than 7-fold that in the control condition were analysed using Gene Ontology enrichment analysis (Ashburner et al., 2000; The Gene Ontology, 2019), searching for enriched biological processes.

RNAPII binding variation between conditions was calculated using z-scores, which were calculated as follows: $Zscore=\frac{\left( KD -WT \right)}{\sqrt{\frac{\left( KD +WT \right)}{2}}}$ (i.e. difference weighted by mean signal), which transforms the distribution of variations into a normal distribution, allowing for better statistical interpretation of the variations.

Scatter plots, box plots, violin plots, pie charts and bar plots were created using either basic R plotting functions or ggplot2 ([https://ggplot2.tidyverse.org](https://ggplot2.tidyverse.org/)) functions. Average binding profiles of proteins across genomic features of interest were generated using seqplots’ (DOI: [10.18129/B9.bioc.seqplots](https://doi.org/doi:10.18129/B9.bioc.seqplots)) plotAverage function. Heatmaps on genomic features were created using genomation’s (DOI: [10.18129/B9.bioc.genomation](https://doi.org/doi:10.18129/B9.bioc.genomation)) gridHeat function, using the profile matrices generated by DeepTools. Gene set enrichment analysis was performed using fgsea (DOI: [10.18129/B9.bioc.fgsea](https://doi.org/doi:10.18129/B9.bioc.fgsea)), and all enrichment plots were created using the plotEnrichment function of the package. Decile matrices were created using color2D matplot from the plotrix package (https://www.rdocumentation.org/packages/plotrix/versions/3.7-7). Genes ranked by z-score of RNAPII variation were split into deciles, for both shMRE11 vs shCon and shNBS1 vs shCon. Then, for each pair of deciles, the number of genes in the intersection of those two deciles was saved into a matrix, creating a 10x10 matrix of integers. The number of genes in each intersection was tested for significance by hypergeometric test, creating a 10x10 matrix of p-values. The genes at each intersection were also tested by hypergeometric test for significant enrichment in either top MRE11 or NBS1 binding levels (decile 1 of genes ranked by MRE11 or NBS1 binding).

For analysis of whole genome sequencing data, following processing by Annovar (Wang et al., 2010) on the sequencing platform, SNP location was then overlapped with gene bodies using Genomic Ranges functions, which yields a quantification per gene.

**ACCESSION NUMBERS**

ChIP-seq data have been deposited at GEO (GSE 143591).
