## Supplementary material for "Chromatin-associated MRN complex protects highly transcribing genes from genomic instability": suppl figs

### Slide 1
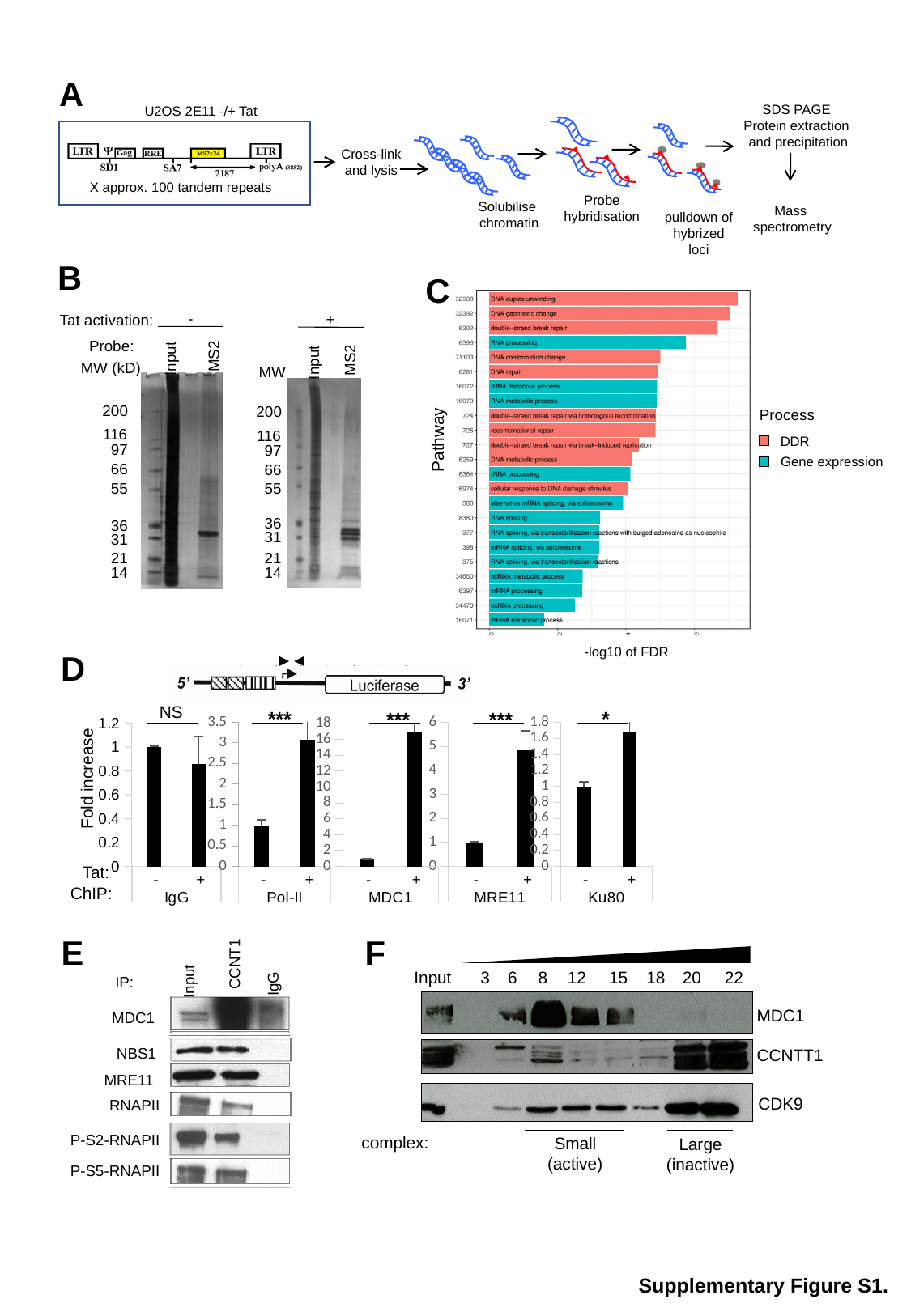

A
SDS PAGE
Protein extraction
 and precipitation
U2OS 2E11 -/+ Tat
Cross-link and lysis
X approx. 100 tandem repeats
 Probe
 hybridisation
Solubilise
chromatin
Mass
spectrometry
pulldown of hybrized
loci
B
C
-
+
Tat activation:
Probe:
MS2
Input
MW (kD)
MS2
Input
MW
200
116
97
66
55
36
31
21
14
200
116
97
66
55
36
31
21
14
Process
Pathway
DDR
Gene expression
-log10 of FDR
D
NS
*
***
***
***
#### Chart
| Category | |
|---|---|
| - | 1.0 |
| + | 0.860063179011531 |
#### Chart
| Category | |
|---|---|
| - | 1.0 |
| + | 3.07958291417164 |
#### Chart
| Category | |
|---|---|
| - | 1.0 |
| + | 4.836536859949565 |
#### Chart
| Category | |
|---|---|
| - | 1.0 |
| + | 1.67055070532455 |
#### Chart
| Category | |
|---|---|
| - | 1.0 |
| + | 16.94206329164128 |Fold increase
Tat:
ChIP:
E
CCNT1
IP:
Input
IgG
MDC1
NBS1
MRE11
RNAPII
P-S2-RNAPII
P-S5-RNAPII
F
Input
3
6
8
12
15
18
20
22
MDC1
CCNTT1
CDK9
complex:
Small (active)
Large (inactive)
Supplementary Figure S1.

### Slide 2
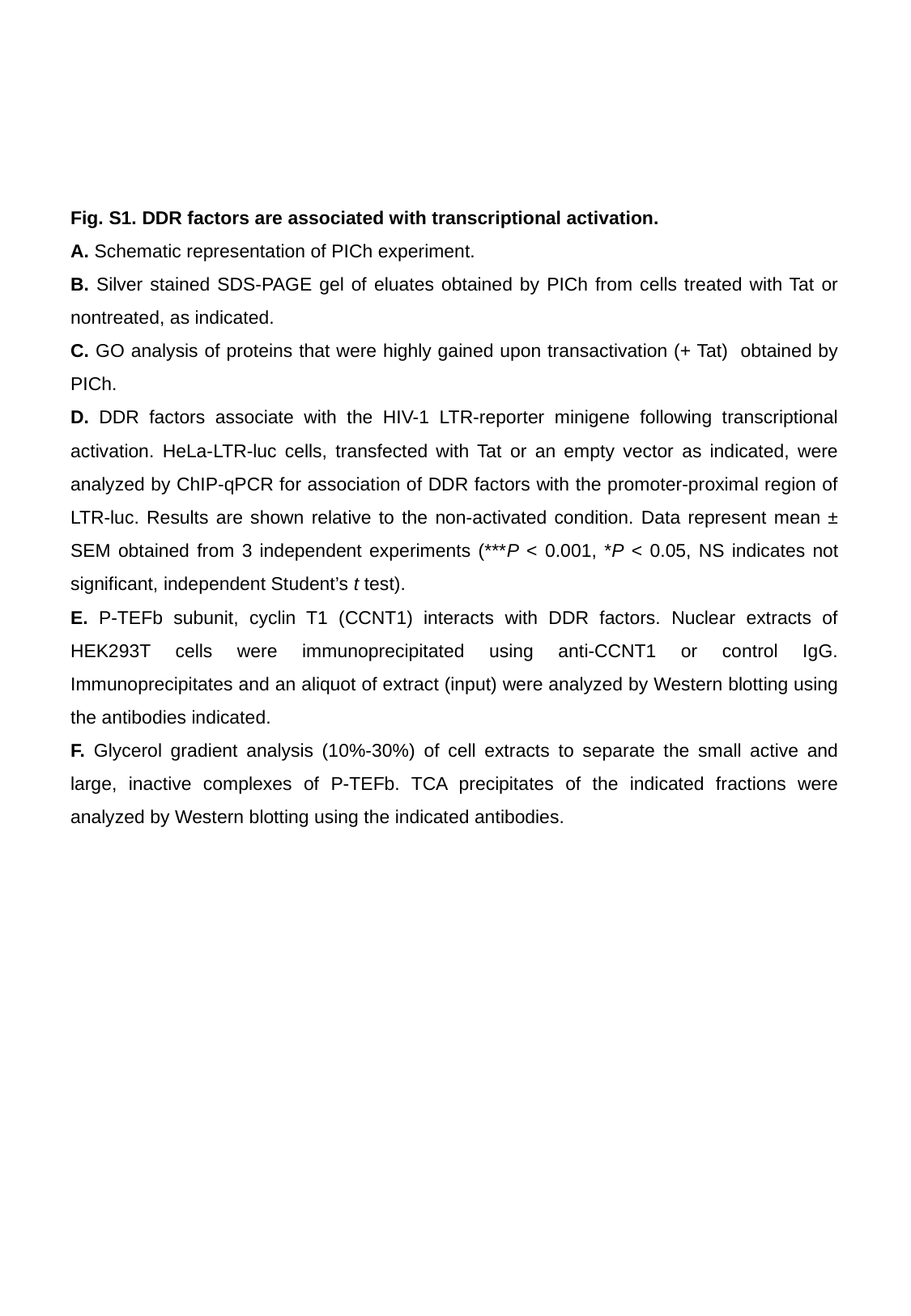

Fig. S1. DDR factors are associated with transcriptional activation.
A. Schematic representation of PICh experiment.
B. Silver stained SDS-PAGE gel of eluates obtained by PICh from cells treated with Tat or nontreated, as indicated.
C. GO analysis of proteins that were highly gained upon transactivation (+ Tat) obtained by PICh.
D. DDR factors associate with the HIV-1 LTR-reporter minigene following transcriptional activation. HeLa-LTR-luc cells, transfected with Tat or an empty vector as indicated, were analyzed by ChIP-qPCR for association of DDR factors with the promoter-proximal region of LTR-luc. Results are shown relative to the non-activated condition. Data represent mean ± SEM obtained from 3 independent experiments (***P < 0.001, *P < 0.05, NS indicates not significant, independent Student’s t test).
E. P-TEFb subunit, cyclin T1 (CCNT1) interacts with DDR factors. Nuclear extracts of HEK293T cells were immunoprecipitated using anti-CCNT1 or control IgG. Immunoprecipitates and an aliquot of extract (input) were analyzed by Western blotting using the antibodies indicated.
F. Glycerol gradient analysis (10%-30%) of cell extracts to separate the small active and large, inactive complexes of P-TEFb. TCA precipitates of the indicated fractions were analyzed by Western blotting using the indicated antibodies.

### Slide 3
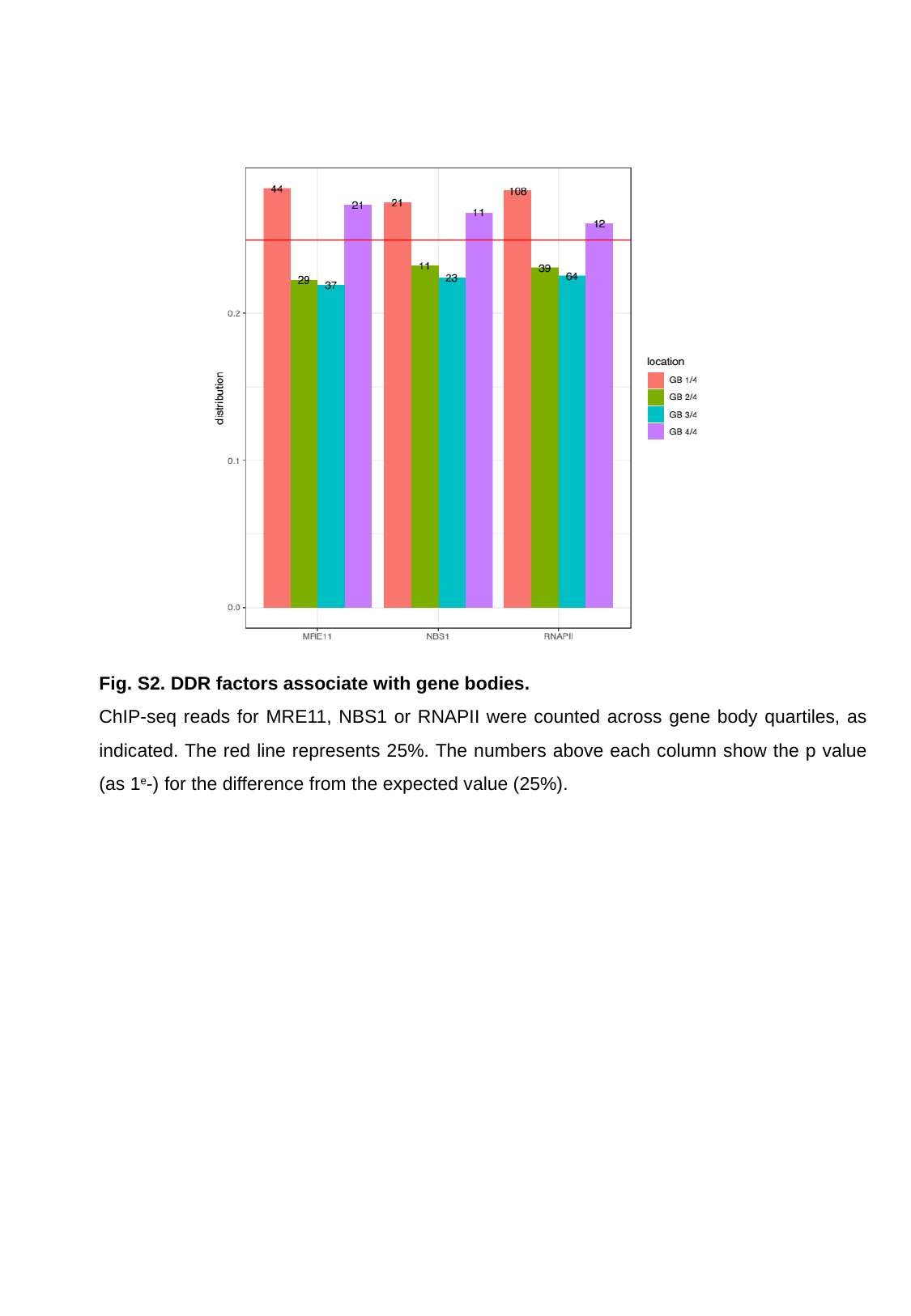

Fig. S2. DDR factors associate with gene bodies.
ChIP-seq reads for MRE11, NBS1 or RNAPII were counted across gene body quartiles, as indicated. The red line represents 25%. The numbers above each column show the p value (as 1^e^-) for the difference from the expected value (25%).

### Slide 4
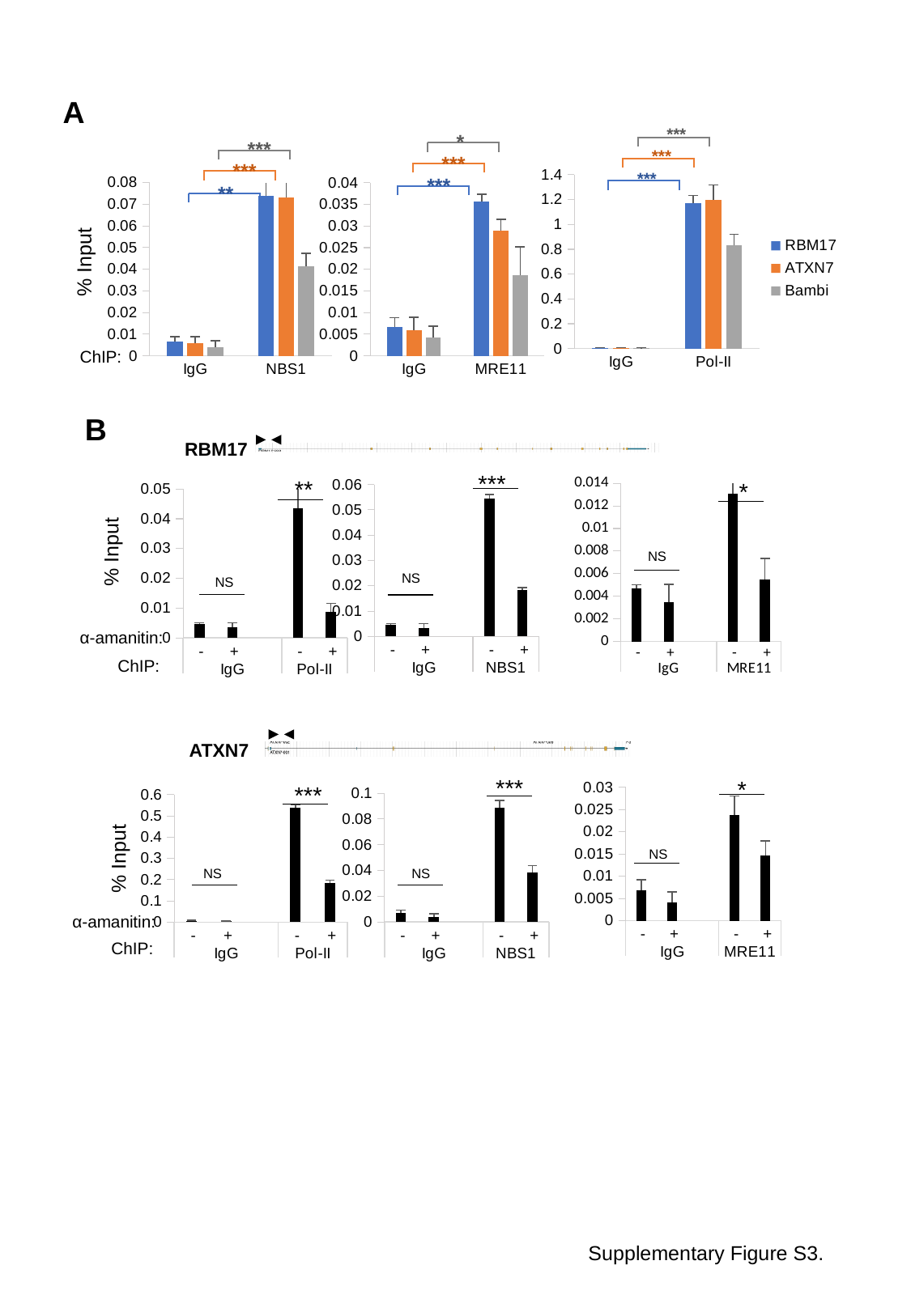

A
***
***
***
*
***
***
***
***
**
#### Chart
| Category | RBM17 | ATXN7 | Bambi |
|---|---|---|---|
| IgG | 0.0066262765201946 | 0.00594003949978686 | 0.00414819370038155 |
| Pol-II | 1.171011132607495 | 1.196697238460041 | 0.83000819339329 |
#### Chart
| Category | RBM17 | ATXN7 | Bambi |
|---|---|---|---|
| IgG | 0.0066262765201946 | 0.00594003949978686 | 0.00414819370038155 |
| NBS1 | 0.073891968149419 | 0.0732406868007033 | 0.0414412417299711 |
#### Chart
| Category | RBM17 | ATXN7 | Bambi |
|---|---|---|---|
| IgG | 0.0066262765201946 | 0.00594003949978686 | 0.00414819370038155 |
| MRE11 | 0.0355948817454738 | 0.0288683126294481 | 0.0186370515710984 |% Input
ChIP:
B
RBM17
***
#### Chart
| Category | RBM17 |
|---|---|
| - | 0.00468809912239941 |
| + | 0.00345962378287444 |
| | None |
| - | 0.0543503700594123 |
| + | 0.0183828654916674 |NS
**
NS
#### Chart
| Category | RBM17 |
|---|---|
| - | 0.00468809912239941 |
| + | 0.00345962378287444 |
| | None |
| - | 0.0436248997563328 |
| + | 0.0087393234422108 |*
#### Chart
| Category | RBM17 |
|---|---|
| - | 0.00468809912239941 |
| + | 0.00345962378287444 |
| | None |
| - | 0.0130531945367779 |
| + | 0.00546822453611521 |NS
% Input
α-amanitin:
ChIP:
ATXN7
***
#### Chart
| Category | ATXN7 |
|---|---|
| - | 0.0067950127758589 |
| + | 0.00405973715866359 |
| | None |
| - | 0.0886245370682729 |
| + | 0.0384648408056481 |NS
*
#### Chart
| Category | ATXN7 |
|---|---|
| - | 0.0067950127758589 |
| + | 0.00405973715866359 |
| | None |
| - | 0.0238311251389182 |
| + | 0.0147021407027846 |NS
***
NS
#### Chart
| Category | ATXN7 |
|---|---|
| - | 0.0067950127758589 |
| + | 0.00405973715866359 |
| | None |
| - | 0.539081154461323 |
| + | 0.185440390475088 |% Input
α-amanitin:
ChIP:
Supplementary Figure S3.

### Slide 5
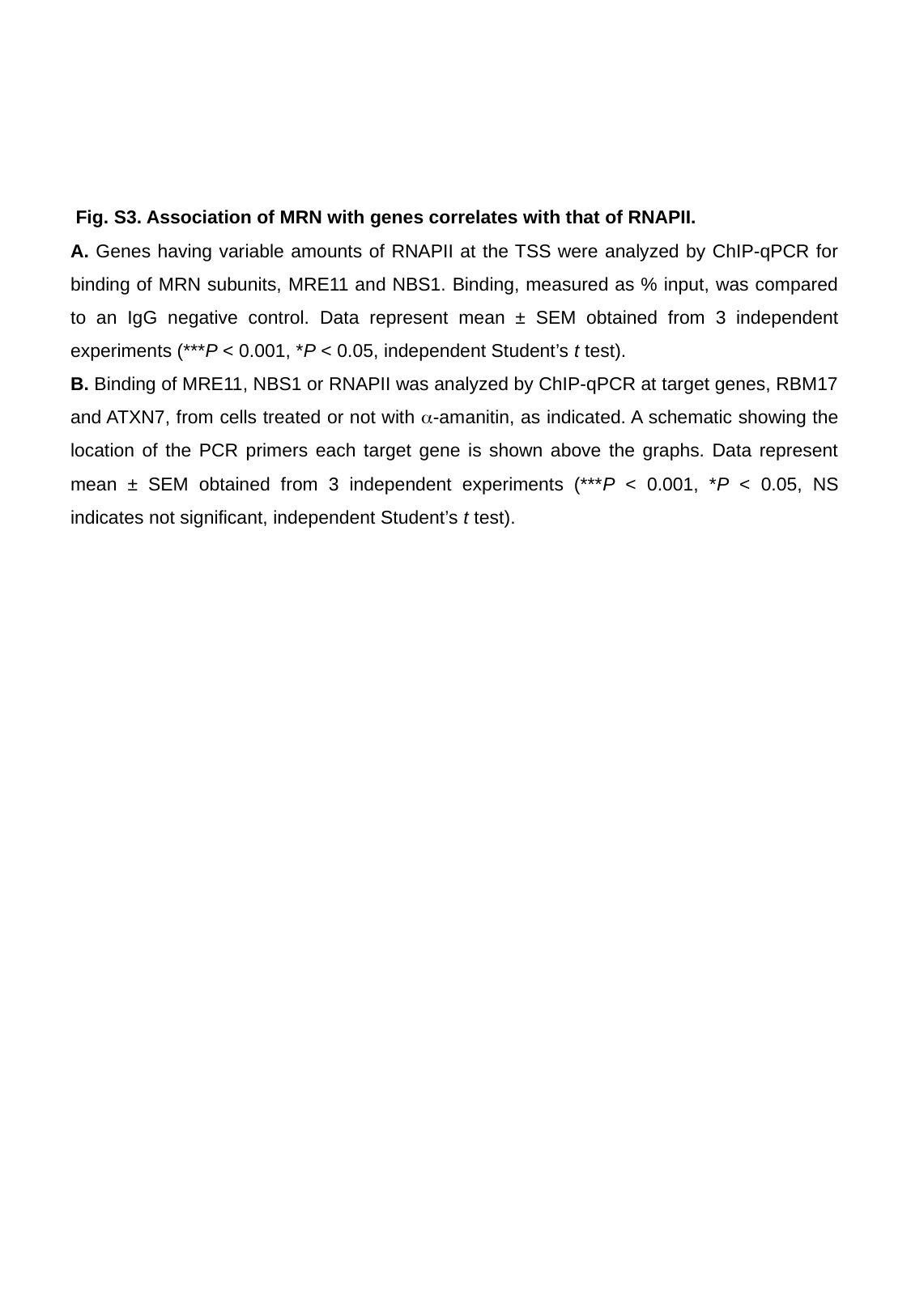

Fig. S3. Association of MRN with genes correlates with that of RNAPII.
A. Genes having variable amounts of RNAPII at the TSS were analyzed by ChIP-qPCR for binding of MRN subunits, MRE11 and NBS1. Binding, measured as % input, was compared to an IgG negative control. Data represent mean ± SEM obtained from 3 independent experiments (***P < 0.001, *P < 0.05, independent Student’s t test).
B. Binding of MRE11, NBS1 or RNAPII was analyzed by ChIP-qPCR at target genes, RBM17 and ATXN7, from cells treated or not with α-amanitin, as indicated. A schematic showing the location of the PCR primers each target gene is shown above the graphs. Data represent mean ± SEM obtained from 3 independent experiments (***P < 0.001, *P < 0.05, NS indicates not significant, independent Student’s t test).

### Slide 6
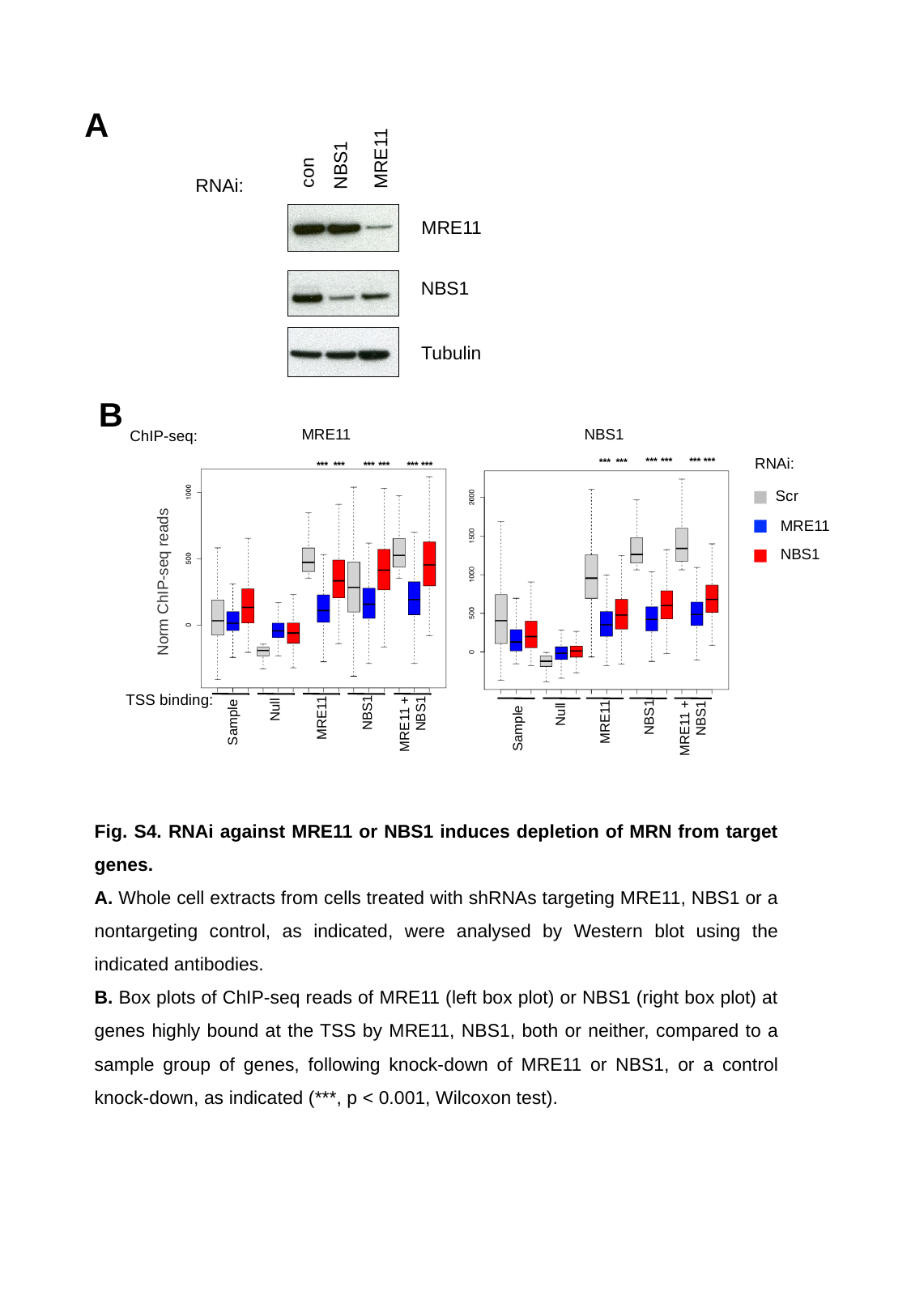

A
MRE11
NBS1
con
RNAi:
MRE11
NBS1
Tubulin
B
NBS1
MRE11
ChIP-seq:
RNAi:
***
***
***
***
***
***
***
***
***
***
***
***
Scr
MRE11
NBS1
Norm ChIP-seq reads
Null
 MRE11 + NBS1
Sample
 NBS1
MRE11
TSS binding:
Null
 MRE11 + NBS1
 NBS1
Sample
MRE11
Fig. S4. RNAi against MRE11 or NBS1 induces depletion of MRN from target genes.
A. Whole cell extracts from cells treated with shRNAs targeting MRE11, NBS1 or a nontargeting control, as indicated, were analysed by Western blot using the indicated antibodies.
B. Box plots of ChIP-seq reads of MRE11 (left box plot) or NBS1 (right box plot) at genes highly bound at the TSS by MRE11, NBS1, both or neither, compared to a sample group of genes, following knock-down of MRE11 or NBS1, or a control knock-down, as indicated (***, p < 0.001, Wilcoxon test).

### Slide 7
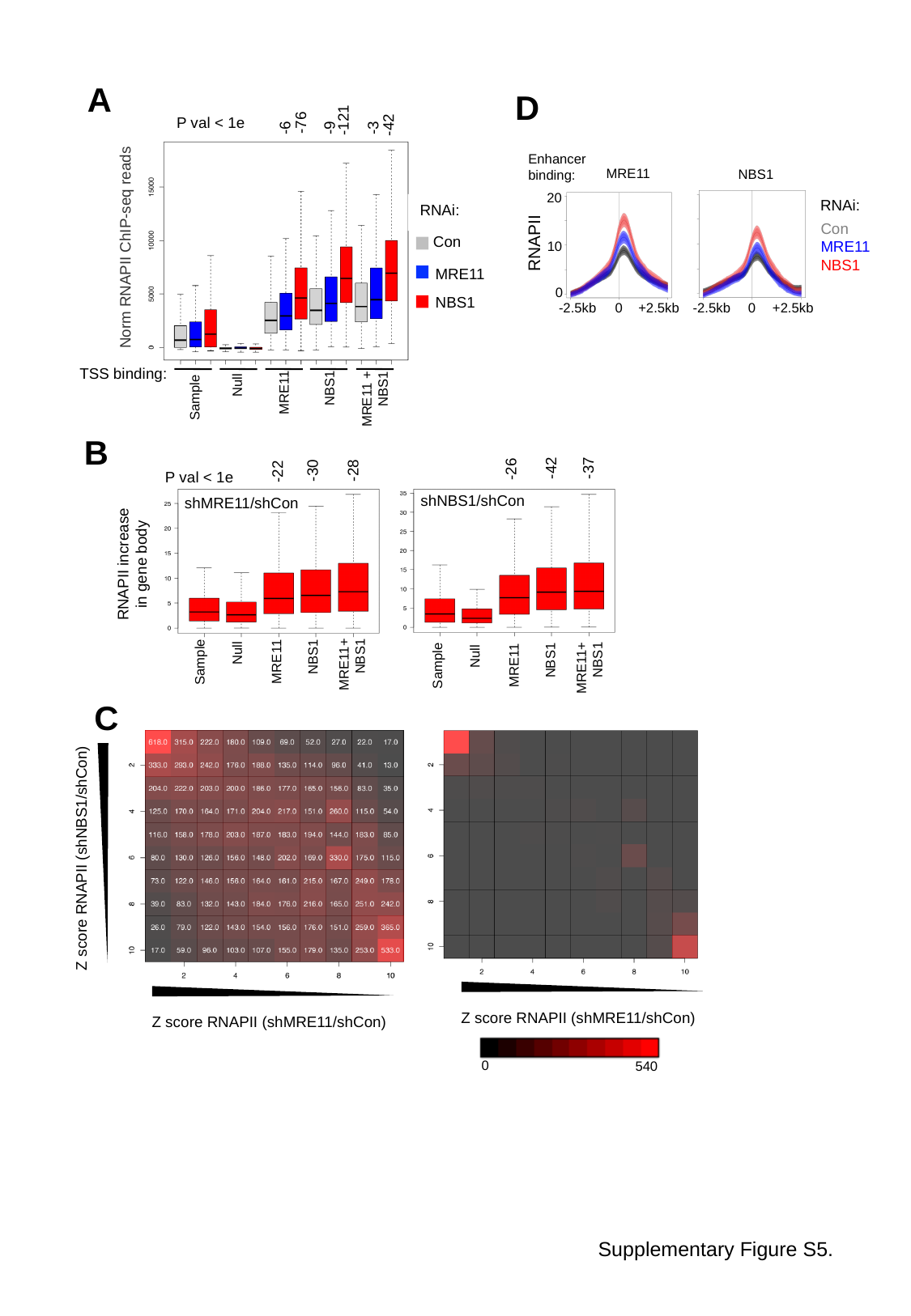

A
D
-121
-76
P val < 1e
-42
-6
-9
-3
Enhancer binding:
MRE11
NBS1
20
RNAi:
RNAi:
Con
MRE11
NBS1
Con
RNAPII
10
Norm RNAPII ChIP-seq reads
MRE11
0
NBS1
-2.5kb
0
+2.5kb
-2.5kb
0
+2.5kb
TSS binding:
Null
 MRE11 + NBS1
Sample
 NBS1
MRE11
B
-42
-37
-26
-30
-28
-22
P val < 1e
shNBS1/shCon
shMRE11/shCon
RNAPII increase in gene body
NBS1
MRE11+NBS1
MRE11
Sample
Null
NBS1
MRE11+NBS1
MRE11
Sample
Null
C
Z score RNAPII (shNBS1/shCon)
Z score RNAPII (shMRE11/shCon)
Z score RNAPII (shMRE11/shCon)
0
540
Supplementary Figure S5.

### Slide 8
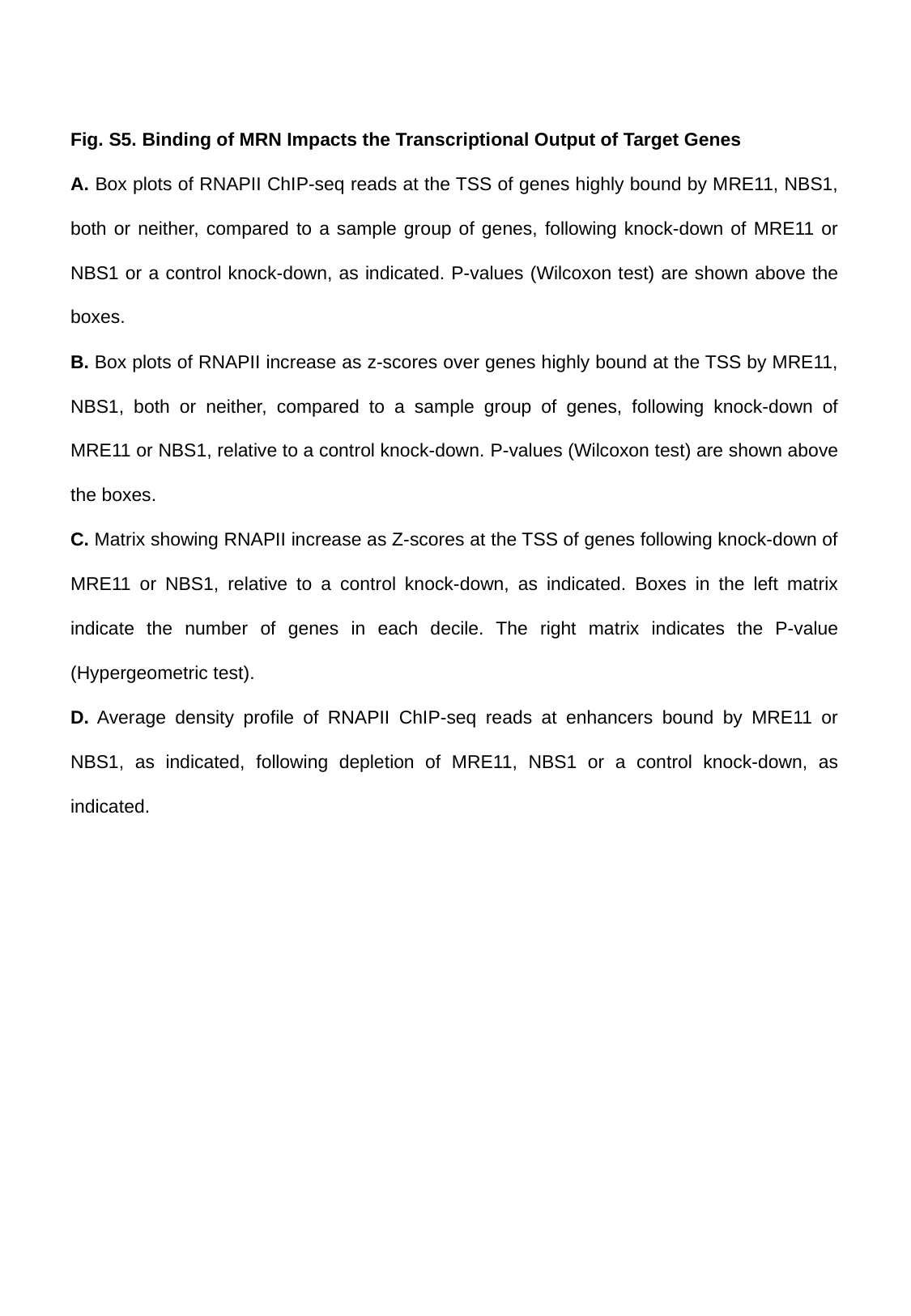

Fig. S5. Binding of MRN Impacts the Transcriptional Output of Target Genes
A. Box plots of RNAPII ChIP-seq reads at the TSS of genes highly bound by MRE11, NBS1, both or neither, compared to a sample group of genes, following knock-down of MRE11 or NBS1 or a control knock-down, as indicated. P-values (Wilcoxon test) are shown above the boxes.
B. Box plots of RNAPII increase as z-scores over genes highly bound at the TSS by MRE11, NBS1, both or neither, compared to a sample group of genes, following knock-down of MRE11 or NBS1, relative to a control knock-down. P-values (Wilcoxon test) are shown above the boxes.
C. Matrix showing RNAPII increase as Z-scores at the TSS of genes following knock-down of MRE11 or NBS1, relative to a control knock-down, as indicated. Boxes in the left matrix indicate the number of genes in each decile. The right matrix indicates the P-value (Hypergeometric test).
D. Average density profile of RNAPII ChIP-seq reads at enhancers bound by MRE11 or NBS1, as indicated, following depletion of MRE11, NBS1 or a control knock-down, as indicated.
