## Supplementary material for "Chromatin-associated MRN complex protects highly transcribing genes from genomic instability": suppl tables

**Table S1.** Identification of proteins whose association with the HIV-1 minigene was significantly changed in the presence Tat transcriptional activator compared to the nontreated sample.

| UniProt ID | logFC | UniProt ID | logFC | UniProt ID | logFC | UniProt ID | logFC |
| --- | --- | --- | --- | --- | --- | --- | --- |
| RL1D1_HUMAN | 3,61 | PUF60_HUMAN | 2,08 | IMA2_HUMAN | 1,79 | PDIA4_HUMAN | 1,61 |
| HP1B3_HUMAN | 3,47 | G6PI_HUMAN | 2,08 | MECP2_HUMAN | 1,79 | SPAS2_HUMAN | 1,61 |
| RPA34_HUMAN | 3,33 | MOGS_HUMAN | 2,08 | FUBP1_HUMAN | 1,79 | SYK_HUMAN | 1,61 |
| MDC1_HUMAN | 3,28 | CALX_HUMAN | 2,08 | RFC4_HUMAN | 1,79 | SSRP1_HUMAN | 1,61 |
| XRCC5_HUMAN | 3,22 | HSP72_HUMAN | 2,08 | CCD47_HUMAN | 1,79 | FUS_HUMAN | 1,61 |
| HNRPM_HUMAN | 3,22 | MCM7_HUMAN | 2,08 | SYDC_HUMAN | 1,79 | CX067_HUMAN | 1,61 |
| TKT_HUMAN | 3,18 | PA2G4_HUMAN | 2,08 | COX2_HUMAN | 1,79 | HAT1_HUMAN | 1,61 |
| AHNK_HUMAN | 3,09 | GRP75_HUMAN | 2,05 | HSP7C_HUMAN | 1,70 | AATM_HUMAN | 1,61 |
| NOP56_HUMAN | 3,09 | LDHB_HUMAN | 2,01 | DHX9_HUMAN | 1,70 | VRK1_HUMAN | 1,61 |
| MCM6_HUMAN | 2,94 | ENPL_HUMAN | 1,95 | ATPA_HUMAN | 1,70 | NCLN_HUMAN | 1,61 |
| H90B3_HUMAN | 2,94 | SYEP_HUMAN | 1,95 | NPM_HUMAN | 1,70 | RBBP4_HUMAN | 1,61 |
| HS90A_HUMAN | 2,83 | BOP1_HUMAN | 1,95 | HSP71_HUMAN | 1,67 | QCR2_HUMAN | 1,61 |
| XRCC6_HUMAN | 2,77 | MCM4_HUMAN | 1,95 | SYIC_HUMAN | 1,61 | PRS6A_HUMAN | 1,61 |
| MOES_HUMAN | 2,77 | UBP5_HUMAN | 1,95 | ACTN1_HUMAN | 1,61 | HNRDL_HUMAN | 1,61 |
| MCM5_HUMAN | 2,64 | EIF3C_HUMAN | 1,95 | PSIP1_HUMAN | 1,61 | ROA1_HUMAN | 1,61 |
| HNRPR_HUMAN | 2,64 | SRSF3_HUMAN | 1,95 | HMGB1_HUMAN | 1,61 | HMGA2_HUMAN | 1,61 |
| DDX17_HUMAN | 2,64 | IMB1_HUMAN | 1,95 | RPA1_HUMAN | 1,61 | RS15_HUMAN | 1,61 |
| PHB2_HUMAN | 2,64 | NAT10_HUMAN | 1,95 | RRP5_HUMAN | 1,61 | DEK_HUMAN | 1,58 |
| TFR1_HUMAN | 2,56 | TSR1_HUMAN | 1,95 | HEAT1_HUMAN | 1,61 | HNRPD_HUMAN | 1,54 |
| DDX21_HUMAN | 2,48 | ILF2_HUMAN | 1,95 | TRIPC_HUMAN | 1,61 | NUCL_HUMAN | 1,53 |
| TIF1B_HUMAN | 2,48 | DDX18_HUMAN | 1,95 | E7ETY2_HUMAN | 1,61 | P5CS_HUMAN | 1,50 |
| DDX27_HUMAN | 2,48 | LKHA4_HUMAN | 1,95 | FND3B_HUMAN | 1,61 | HSP74_HUMAN | 1,50 |
| RECQ1_HUMAN | 2,48 | ZN512_HUMAN | 1,95 | STT3B_HUMAN | 1,61 | HDGF_HUMAN | 1,50 |
| LMNB2_HUMAN | 2,47 | DDX1_HUMAN | 1,95 | LRRF1_HUMAN | 1,61 | RS2_HUMAN | 1,47 |
| SFPQ_HUMAN | 2,40 | NOG1_HUMAN | 1,95 | SMHD1_HUMAN | 1,61 | BAF_HUMAN | 1,47 |
| KI67_HUMAN | 2,30 | FBRL_HUMAN | 1,95 | NOL6_HUMAN | 1,61 | GRP78_HUMAN | 1,46 |
| RRMJ3_HUMAN | 2,30 | TGM2_HUMAN | 1,95 | NOMO1_HUMAN | 1,61 | MCM2_HUMAN | 1,45 |
| RBM39_HUMAN | 2,30 | ABCE1_HUMAN | 1,95 | ACLY_HUMAN | 1,61 | LDHA_HUMAN | 1,45 |
| DDX3X_HUMAN | 2,30 | EIF3L_HUMAN | 1,95 | SMCA1_HUMAN | 1,61 | LAP2A_HUMAN | 1,42 |
| KV309_HUMAN | 2,30 | DNJC9_HUMAN | 1,95 | WDR12_HUMAN | 1,61 | SP16H_HUMAN | 1,39 |
| TCPB_HUMAN | 2,30 | LA_HUMAN | 1,95 | MYO1C_HUMAN | 1,61 | HS90B_HUMAN | 1,39 |
| DDX5_HUMAN | 2,30 | LAP2B_HUMAN | 1,95 | SUN2_HUMAN | 1,61 | ALDOC_HUMAN | 1,39 |
| MCM3_HUMAN | 2,25 | THOC4_HUMAN | 1,95 | DCTN2_HUMAN | 1,61 | TOP1M_HUMAN | 1,39 |
| ESYT1_HUMAN | 2,20 | NOP2_HUMAN | 1,92 | CTNA1_HUMAN | 1,61 | H2AW_HUMAN | 1,39 |
| DDB1_HUMAN | 2,20 | LYRIC_HUMAN | 1,90 | STAT1_HUMAN | 1,61 | RCC1_HUMAN | 1,39 |
| IMMT_HUMAN | 2,20 | NUMA1_HUMAN | 1,87 | HDGR2_HUMAN | 1,61 | APEX1_HUMAN | 1,39 |
| STIP1_HUMAN | 2,20 | UBA1_HUMAN | 1,87 | UHRF1_HUMAN | 1,61 | CHD4_HUMAN | 1,39 |
| ASNS_HUMAN | 2,20 | TOP2A_HUMAN | 1,79 | UBP2L_HUMAN | 1,61 | SPT6H_HUMAN | 1,39 |
| MK67I_HUMAN | 2,20 | FLNB_HUMAN | 1,79 | HNRPF_HUMAN | 1,61 | KIF4A_HUMAN | 1,39 |
| TM214_HUMAN | 2,20 | GCN1L_HUMAN | 1,79 | CMC2_HUMAN | 1,61 | MSH6_HUMAN | 1,39 |
| YBOX1_HUMAN | 2,20 | SYLC_HUMAN | 1,79 | PUR9_HUMAN | 1,61 | MBB1A_HUMAN | 1,39 |
| GUAA_HUMAN | 2,20 | MAGD2_HUMAN | 1,79 | NSUN2_HUMAN | 1,61 | BAZ2A_HUMAN | 1,39 |
| HNRPQ_HUMAN | 2,20 | ACTN4_HUMAN | 1,79 | EWS_HUMAN | 1,61 | NOLC1_HUMAN | 1,39 |
| HNRCL_HUMAN | 2,20 | PABP1_HUMAN | 1,79 | VPS35_HUMAN | 1,61 | ASPH_HUMAN | 1,39 |
| LMNB1_HUMAN | 2,13 | ITB1_HUMAN | 1,79 | FUBP2_HUMAN | 1,61 | TTF1_HUMAN | 1,39 |
| CKAP4_HUMAN | 2,12 | DLDH_HUMAN | 1,79 | RCC2_HUMAN | 1,61 | FXR1_HUMAN | 1,39 |
| BAZ1B_HUMAN | 2,08 | ILF3_HUMAN | 1,79 | D6RBZ0_HUMAN | 1,61 | PWP2_HUMAN | 1,39 |
| LPPRC_HUMAN | 2,08 | DHX15_HUMAN | 1,79 | DPYL3_HUMAN | 1,61 | XPC_HUMAN | 1,39 |
| TBL3_HUMAN | 2,08 | SYQ_HUMAN | 1,79 | GNL3_HUMAN | 1,61 | WDR36_HUMAN | 1,39 |

| UniProt ID | logFC | UniProt ID | logFC | UniProt ID | logFC | UniProt ID | logFC |
| --- | --- | --- | --- | --- | --- | --- | --- |
| MAN1_HUMAN | 1,39 | UBC9_HUMAN | 1,39 | EIF3B_HUMAN | 0,98 | CPNE3_HUMAN | -0,92 |
| COPG1_HUMAN | 1,39 | FLNC_HUMAN | 1,36 | TCP4_HUMAN | 0,98 | RL15_HUMAN | -0,92 |
| ACSL3_HUMAN | 1,39 | COF1_HUMAN | 1,34 | ARF1_HUMAN | 0,98 | ACTB_HUMAN | -0,99 |
| XPO1_HUMAN | 1,39 | MAP1B_HUMAN | 1,34 | TBB1_HUMAN | 0,96 | CATA_HUMAN | -1,10 |
| MVP_HUMAN | 1,39 | LMNA_HUMAN | 1,32 | HNRH1_HUMAN | 0,96 | NCRP1_HUMAN | -1,10 |
| ABCF1_HUMAN | 1,39 | CLIC1_HUMAN | 1,30 | FEN1_HUMAN | 0,96 | ECHB_HUMAN | -1,39 |
| XRN2_HUMAN | 1,39 | SRRT_HUMAN | 1,25 | HMGB3_HUMAN | 0,96 | TGM5_HUMAN | -1,39 |
| U5S1_HUMAN | 1,39 | SMCA5_HUMAN | 1,25 | HMGB2_HUMAN | 0,96 | ADRO_HUMAN | -1,39 |
| NCKP1_HUMAN | 1,39 | RS6_HUMAN | 1,25 | EZRI_HUMAN | 0,94 | DHCR7_HUMAN | -1,39 |
| MATR3_HUMAN | 1,39 | TBA3C_HUMAN | 1,25 | RSSA_HUMAN | 0,94 | DPM1_HUMAN | -1,39 |
| ELYS_HUMAN | 1,39 | DBPA_HUMAN | 1,25 | H2AY_HUMAN | 0,93 | BAP31_HUMAN | -1,39 |
| DDX56_HUMAN | 1,39 | SERPH_HUMAN | 1,25 | PTN1_HUMAN | 0,92 | RU2B_HUMAN | -1,39 |
| TBR1_HUMAN | 1,39 | HS105_HUMAN | 1,25 | TBB4B_HUMAN | 0,92 | RL35A_HUMAN | -1,39 |
| PSMD2_HUMAN | 1,39 | LASP1_HUMAN | 1,25 | BUB3_HUMAN | 0,92 | RL34_HUMAN | -1,39 |
| NOL11_HUMAN | 1,39 | RL6_HUMAN | 1,25 | VDAC2_HUMAN | 0,92 | SMD1_HUMAN | -1,39 |
| ECHA_HUMAN | 1,39 | HNRPU_HUMAN | 1,24 | AL7A1_HUMAN | 0,92 | PYC_HUMAN | -1,61 |
| PESC_HUMAN | 1,39 | TBB5_HUMAN | 1,22 | PGAM1_HUMAN | 0,92 | KV201_HUMAN | -1,61 |
| SRPR_HUMAN | 1,39 | NOP58_HUMAN | 1,22 | RS26_HUMAN | 0,92 | 2AAA_HUMAN | -1,61 |
| SYRC_HUMAN | 1,39 | FLNA_HUMAN | 1,21 | NACA_HUMAN | 0,92 | GP179_HUMAN | -1,61 |
| E2AK2_HUMAN | 1,39 | MYH9_HUMAN | 1,20 | RL8_HUMAN | 0,92 | RUVB2_HUMAN | -1,61 |
| NPL4_HUMAN | 1,39 | TLN1_HUMAN | 1,20 | DDX6_HUMAN | 0,92 | H14_HUMAN | -1,61 |
| TCPE_HUMAN | 1,39 | RL13_HUMAN | 1,20 | CNDP2_HUMAN | 0,92 | FABP5_HUMAN | -1,70 |
| GFPT1_HUMAN | 1,39 | DDX47_HUMAN | 1,20 | GDIR1_HUMAN | 0,92 | ACTBL_HUMAN | -1,79 |
| RFA1_HUMAN | 1,39 | SNUT1_HUMAN | 1,20 | RS3A_HUMAN | 0,92 | POF1B_HUMAN | -1,79 |
| STRAP_HUMAN | 1,39 | RL19_HUMAN | 1,20 | RS29_HUMAN | 0,92 | DHRS2_HUMAN | -1,79 |
| ROA3_HUMAN | 1,39 | RL7_HUMAN | 1,20 | CALM_HUMAN | 0,92 | ACTBM_HUMAN | -1,95 |
| ELAV1_HUMAN | 1,39 | CG050_HUMAN | 1,20 | AK1A1_HUMAN | 0,92 | TRFL_HUMAN | -2,20 |
| PAL4A_HUMAN | 1,39 | PARP1_HUMAN | 1,19 | ANXA5_HUMAN | 0,92 | HUTH_HUMAN | -2,40 |
| G3PT_HUMAN | 1,39 | RBMX_HUMAN | 1,18 | RHOA_HUMAN | 0,92 | FRPD4_HUMAN | -2,48 |
| CAP1_HUMAN | 1,39 | H90B2_HUMAN | 1,16 | RS23_HUMAN | 0,92 |  |  |
| NP1L1_HUMAN | 1,39 | HGB1A_HUMAN | 1,15 | PLEC_HUMAN | 0,91 |  |  |
| IMP3_HUMAN | 1,39 | ROA2_HUMAN | 1,15 | VIME_HUMAN | 0,91 |  |  |
| ACTZ_HUMAN | 1,39 | CLH1_HUMAN | 1,10 | ENOA_HUMAN | 0,90 |  |  |
| BCCIP_HUMAN | 1,39 | TOP1_HUMAN | 1,10 | RS18_HUMAN | 0,88 |  |  |
| PLIN3_HUMAN | 1,39 | LBR_HUMAN | 1,10 | PRKDC_HUMAN | 0,86 |  |  |
| CDC37_HUMAN | 1,39 | PEBP1_HUMAN | 1,10 | RL10_HUMAN | 0,85 |  |  |
| METK2_HUMAN | 1,39 | RPN2_HUMAN | 1,10 | TBB3_HUMAN | 0,85 |  |  |
| EIF3E_HUMAN | 1,39 | G3BP1_HUMAN | 1,10 | SERA_HUMAN | 0,85 |  |  |
| FCL_HUMAN | 1,39 | TPIS_HUMAN | 1,10 | RBM8A_HUMAN | 0,85 |  |  |
| EBP2_HUMAN | 1,39 | RL18_HUMAN | 1,10 | TBB4A_HUMAN | 0,83 |  |  |
| DHSO_HUMAN | 1,39 | DNJB1_HUMAN | 1,10 | RL3_HUMAN | 0,83 |  |  |
| BIEA_HUMAN | 1,39 | IF2A_HUMAN | 1,10 | H2B1A_HUMAN | 0,82 |  |  |
| RLA0L_HUMAN | 1,39 | HNRH3_HUMAN | 1,10 | PUR6_HUMAN | 0,81 |  |  |
| H2B2C_HUMAN | 1,39 | H2AV_HUMAN | 1,10 | CBX1_HUMAN | 0,79 |  |  |
| RANG_HUMAN | 1,39 | HMGN2_HUMAN | 1,10 | ATPB_HUMAN | 0,79 |  |  |
| NTPCR_HUMAN | 1,39 | GBLP_HUMAN | 1,06 | PHB_HUMAN | 0,79 |  |  |
| HTR5A_HUMAN | 1,39 | DKC1_HUMAN | 1,03 | LRC59_HUMAN | 0,77 |  |  |
| DUT_HUMAN | 1,39 | PDIA6_HUMAN | 1,03 | CDSN_HUMAN | 0,77 |  |  |
| SKP1_HUMAN | 1,39 | CBX3_HUMAN | 1,03 | RS3_HUMAN | 0,74 |  |  |
| PPAC_HUMAN | 1,39 | HNRPK_HUMAN | 1,02 | RPN1_HUMAN | 0,73 |  |  |
| DUS3_HUMAN | 1,39 | RS8_HUMAN | 1,01 | RS9_HUMAN | -0,81 |  |  |
| ARPC4_HUMAN | 1,39 | STT3A_HUMAN | 0,98 | ZCH23_HUMAN | -0,85 |  |  |
| TMEDA_HUMAN | 1,39 | PPM1G_HUMAN | 0,98 | PRDX2_HUMAN | -0,92 |  |  |

**Table S2.** Identification of proteins interacting with endogenous MDC1 by tandem affinity purification and mass spectrometry. Results are shown as LogFE, (log fold enrichment) of the number of peptides found in MDC1 sample compared to the control sample.

| ID | logFE | ID | logFE | ID | logFE | ID | logFE | ID | logFE |
| --- | --- | --- | --- | --- | --- | --- | --- | --- | --- |
| MDC1 | 4,68 | RPL6 | 1,79 | SRRM1 | 1,39 | ZNF326 | 1,10 | RPL19 | 0,69 |
| RAD50 | 4,26 | SRSF1 | 1,79 | RBM8A | 1,39 | SNRPD3 | 1,10 | GRWD1 | 0,69 |
| CD2AP | 3,69 | FAM175A | 1,79 | DDX1 | 1,39 | PSMA5 | 1,10 | ALDOC | 0,69 |
| NBN | 3,56 | USP28 | 1,79 | ADH5 | 1,39 | TUBAL3 | 1,10 | SRSF5 | 0,69 |
| MRE11A | 3,47 | SLTM | 1,79 | NUMA1 | 1,39 | HSP90AA5P | 1,10 | NEFM | 0,69 |
| TRIP12 | 3,26 | RPS16 | 1,79 | NCOA5 | 1,39 | HMGB2 | 1,10 | RPL34 | 0,69 |
| ANAPC1 | 3,22 | KPNA2 | 1,79 | SNRNP40 | 1,39 | TRA2A | 1,10 | NUCKS1 | 0,69 |
| SH3KBP1 | 3,14 | PPP1CA | 1,79 | HNRNPUL2 | 1,39 | RBM17 | 1,10 | YBX3 | 0,69 |
| PRPF8 | 3,09 | BRE | 1,79 | THRAP3 | 1,39 | ACTN1 | 1,10 | PRPS1 | 0,69 |
| TEX10 | 3,04 | PNP | 1,79 | SNRPA1 | 1,39 | RBBP4 | 1,10 | PREP | 0,69 |
| TP53BP1 | 3,00 | ZFR | 1,79 | RPS13 | 1,39 | GAR1 | 1,10 | CCT7 | 0,69 |
| PELP1 | 2,89 | HNRNPH3 | 1,79 | NME1 | 1,39 | AMOTL1 | 1,10 | HIST1H2AB | 0,69 |
| SF3B1 | 2,89 | XRN2 | 1,79 | ILF3 | 1,34 | GLO1 | 1,10 | RBM39 | 0,69 |
| SNRNP200 | 2,89 | PABPC1 | 1,79 | ILF2 | 1,32 | EIF4A1 | 1,10 | PDHB | 0,69 |
| YLPM1 | 2,77 | PARP1 | 1,79 | NPM1 | 1,30 | HSPH1 | 1,10 | HIST1H2BB | 0,69 |
| CDC27 | 2,77 | RBM6 | 1,79 | HNRNPL | 1,25 | TOP1 | 1,10 | DNAJA1 | 0,69 |
| CDC16 | 2,64 | RPL7 | 1,79 | ELAVL1 | 1,25 | PRMT5 | 1,10 | PRSS1 | 0,69 |
| LAS1L | 2,64 | RPL4 | 1,79 | XRCC6 | 1,25 | MAGOHB | 1,10 | ACLY | 0,69 |
| DDX21 | 2,64 | HNRNPR | 1,61 | CELSR3 | 1,10 | SLC25A5 | 1,10 | MFAP1 | 0,69 |
| ANAPC5 | 2,64 | SYNCRIP | 1,61 | ANP32E | 1,10 | ZNF638 | 1,10 | RPS26 | 0,69 |
| ANAPC7 | 2,64 | RCN2 | 1,61 | STAU1 | 1,10 | RPL24 | 1,10 | TUBA1B | 0,69 |
| ADAR | 2,64 | HNRNPD | 1,61 | HP1BP3 | 1,10 | RPL27 | 1,10 | KHDRBS1 | 0,69 |
| AMOT | 2,64 | LTA4H | 1,61 | SRSF7 | 1,10 | HSP90AB2P | 1,10 | RPL10 | 0,69 |
| ANAPC4 | 2,56 | HNRNPH2 | 1,61 | PSMA2 | 1,10 | RPL8 | 1,10 | ACAT1 | 0,69 |
| SRRM2 | 2,56 | RBM14 | 1,61 | YWHAZ | 1,10 | ATIC | 1,10 | DDB1 | 0,69 |
| HNRNPM | 2,56 | SNRPD2 | 1,61 | RPLP0 | 1,10 | PCNA | 1,10 | PRPF4B | 0,69 |
| HNRNPC | 2,48 | PRRC2A | 1,61 | SAP18 | 1,10 | HSPA5 | 1,05 | CRNKL1 | 0,69 |
| CAPZA1 | 2,48 | DKC1 | 1,61 | PGD | 1,10 | DHX15 | 1,03 | SRSF3 | 0,69 |
| IGF2BP1 | 2,48 | IGF2BP3 | 1,61 | CDC5L | 1,10 | CAPRIN1 | 0,98 | NACA | 0,69 |
| UIMC1 | 2,40 | RAN | 1,61 | SRSF10 | 1,10 | PSAT1 | 0,98 | APOA1 | 0,69 |
| SENP3 | 2,40 | SRSF4 | 1,61 | RNPS1 | 1,10 | SF3B2 | 0,98 | PSMB4 | 0,69 |
| RALY | 2,30 | HNRNPCL3 | 1,50 | PTBP1 | 1,10 | HNRNPUL1 | 0,96 | RPL31 | 0,69 |
| EFTUD2 | 2,30 | PDIA3 | 1,39 | RPLP0P6 | 1,10 | RPL7A | 0,92 | RPS17L | 0,69 |
| BRCC3 | 2,30 | MTHFD1 | 1,39 | EEF1B2 | 1,10 | ALYREF | 0,92 | PRPF6 | 0,69 |
| MATR3 | 2,30 | HNRNPK | 1,39 | PSMB3 | 1,10 | HDGF | 0,92 | CWC22 | 0,69 |
| MDN1 | 2,30 | WDR1 | 1,39 | H1FX | 1,10 | SF3A1 | 0,92 | YWHAQ | 0,69 |
| ANAPC2 | 2,30 | HNRNPA1 | 1,39 | EEF1D | 1,10 | GDI2 | 0,92 | CCT6A | 0,69 |
| SF3B3 | 2,25 | GDI1 | 1,39 | NAP1L1 | 1,10 | RPL9 | 0,92 | HSPE1 | 0,69 |
| WDR18 | 2,20 | ETFA | 1,39 | TARDBP | 1,10 | EZR | 0,92 | SHMT2 | 0,69 |
| TP53 | 2,08 | YWHAB | 1,39 | SNW1 | 1,10 | U2SURP | 0,85 | IMPDH2 | 0,69 |
| CAPZB | 2,08 | RPSA | 1,39 | PYGL | 1,10 | GAPDH | 0,85 | GARS | 0,69 |
| RPL3 | 2,08 | CIRBP | 1,39 | PRKRA | 1,10 | RPS4X | 0,85 | ACTN4 | 0,69 |
| CAPZA2 | 1,95 | TRA2B | 1,39 | CLTC | 1,10 | TPI1 | 0,81 | DLD | 0,69 |
| BABAM1 | 1,95 | PRPF19 | 1,39 | SNRPE | 1,10 | ANXA5 | 0,81 | SSRP1 | 0,69 |
| NOL9 | 1,95 | APEX1 | 1,39 | RBM10 | 1,10 | HSPA9 | 0,79 | CCT2 | 0,69 |
| TJP1 | 1,95 | XRCC5 | 1,39 | RPL27A | 1,10 | DIAPH1 | 0,69 | WDR61 | 0,69 |
| CHERP | 1,95 | RPL15 | 1,39 | HPRT1 | 1,10 | KAT2A | 0,69 | RRP1B | 0,69 |
| USP7 | 1,95 | ANAPC10 | 1,39 | PSMB5 | 1,10 | HSBP1 | 0,69 | SF3B4 | 0,69 |
| RSL1D1 | 1,95 | NOP56 | 1,39 | HIST1H4A | 1,10 | DCDC2C | 0,69 | OAT | 0,69 |
| CDC23 | 1,95 | RPS15A | 1,39 | SUB1 | 1,10 | TMC7 | 0,69 | SHMT1 | 0,69 |
| ID | logFE | ID | logFE | ID | logFE | ID | logFE | ID | logFE |
| RPN2 | 0,69 | CWC22 | 0,69 | NHP2 | 0,69 | HDAC1 | 0,69 | CKB | 0,64 |
| CCT3 | 0,69 | YWHAQ | 0,69 | RPS10 | 0,69 | CPNE3 | 0,69 | AHCY | 0,59 |
| C21orf33 | 0,69 | CCT6A | 0,69 | SKIV2L2 | 0,69 | PSMB6 | 0,69 | UBA1 | 0,59 |
| ZCCHC8 | 0,69 | HSPE1 | 0,69 | LUC7L2 | 0,69 | RPS11 | 0,69 | DDX3X | 0,56 |
| RRP9 | 0,69 | SHMT2 | 0,69 | RDX | 0,69 | SUPT16H | 0,69 | SSB | 0,56 |
| RPL37A | 0,69 | IMPDH2 | 0,69 | RNPEP | 0,69 | PCYOX1 | 0,69 | HNRNPAB | 0,56 |
| FUBP3 | 0,69 | GARS | 0,69 | PFAS | 0,69 | CS | 0,69 | CA2 | 0,56 |
| GTPBP4 | 0,69 | ACTN4 | 0,69 | STRBP | 0,69 | SNRPB2 | 0,69 | HSPA1L | 0,51 |
| TJP2 | 0,69 | DLD | 0,69 | APEH | 0,69 | UHRF1 | 0,69 | GTF2I | 0,51 |
| RPL23 | 0,69 | SSRP1 | 0,69 | BCAS2 | 0,69 | CDH7 | 0,69 | DHX9 | 0,51 |
| RPS28 | 0,69 | CCT2 | 0,69 | OXCT1 | 0,69 | ZNF346 | 0,69 | TALDO1 | 0,51 |
| GSTP1 | 0,69 | WDR61 | 0,69 | GRHPR | 0,69 | FADD | 0,69 | PA2G4 | 0,51 |
| RPLP1 | 0,69 | RRP1B | 0,69 | RPL5 | 0,69 | PARP10 | 0,69 | NASP | 0,51 |
| EIF6 | 0,69 | SF3B4 | 0,69 | SMNDC1 | 0,69 | MAD2L1 | 0,69 | SAFB2 | 0,51 |
| GANAB | 0,69 | OAT | 0,69 | HMGB3 | 0,69 | CLIC1 | 0,69 | MDH1 | 0,51 |
| SERBP1 | 0,69 | SHMT1 | 0,69 | NKRF | 0,69 | PLOD2 | 0,69 | DDX5 | 0,49 |
| STMN1 | 0,69 | RPN2 | 0,69 | SNRPB | 0,69 | FAM171A1 | 0,69 | RBMX | 0,47 |
| RBM39 | 0,69 | CCT3 | 0,69 | PPAT | 0,69 | RPL36 | 0,69 | HSPA4 | 0,47 |
| PDHB | 0,69 | C21orf33 | 0,69 | NUDT5 | 0,69 | TRIO | 0,69 | TUBB2A | 0,45 |
| HIST1H2BB | 0,69 | ZCCHC8 | 0,69 | PABPN1 | 0,69 | E4F1 | 0,69 | TKT | 0,41 |
| DNAJA1 | 0,69 | RRP9 | 0,69 | NCBP1 | 0,69 | RSRC1 | 0,69 | ACTB | 0,41 |
| PRSS1 | 0,69 | RPL37A | 0,69 | PRPH | 0,69 | BRF1 | 0,69 | HNRNPDL | 0,41 |
| ACLY | 0,69 | FUBP3 | 0,69 | ARFIP1 | 0,69 | SF1 | 0,69 | RPS2 | 0,41 |
| MFAP1 | 0,69 | GTPBP4 | 0,69 | CPSF1 | 0,69 | ERH | 0,69 | ANP32A | 0,41 |
| RPS26 | 0,69 | TJP2 | 0,69 | FHL1 | 0,69 | GAPDHS | 0,69 | HIST1H1A | 0,41 |
| TUBA1B | 0,69 | RPL23 | 0,69 | GOT1 | 0,69 | YBX1 | 0,69 | YWHAE | 0,41 |
| KHDRBS1 | 0,69 | RPS28 | 0,69 | SYNE1 | 0,69 | EWSR1 | 0,69 | RPL18 | 0,41 |
| RPL10 | 0,69 | GSTP1 | 0,69 | PTMS | 0,69 | GOT2 | 0,69 | RPS14 | 0,41 |
| ACAT1 | 0,69 | RPLP1 | 0,69 | HIST1H2AA | 0,69 | HNRNPF | 0,69 | RANBP1 | 0,41 |
| DDB1 | 0,69 | EIF6 | 0,69 | FARSA | 0,69 | RPL12 | 0,69 | TUBB3 | 0,41 |
| PRPF4B | 0,69 | GANAB | 0,69 | PRMT1 | 0,69 | PGAM1 | 0,69 | NPEPPS | 0,41 |
| CRNKL1 | 0,69 | STMN1 | 0,69 | ESRRA | 0,69 | TUBB4A | 0,69 | CANX | 0,41 |
| SRSF3 | 0,69 | PCBP1 | 0,69 | DDX50 | 0,69 | ACO2 | 0,69 | PYGB | 0,41 |
| NACA | 0,69 | NANS | 0,69 | KHSRP | 0,69 | PPIB | 0,69 | PSMA6 | 0,41 |
| APOA1 | 0,69 | MSN | 0,69 | SNRPGP15 | 0,69 | PHGDH | 0,69 | POTEKP | 0,41 |
| PSMB4 | 0,69 | SPAG17 | 0,69 | RPL13AP3 | 0,69 | IK | 0,69 | PGM1 | 0,41 |
| RPL31 | 0,69 | EIF4A3 | 0,69 | EIF4H | 0,69 | BDP1 | 0,69 |  |  |
| RPS17L | 0,69 | BSG | 0,69 | PRPF6 | 0,69 | HNRNPU | 0,64 |  |  |

**Table S3.** Antibodies used in this study.

| Antibody | Reference | Supplier |
| --- | --- | --- |
| MDC1 (WB, IP and ChIP) | A300-051A | Bethyl laboratories |
| MRE11 (ChIP, ChIP-seq) | A303-998A | Bethyl laboratories |
| NBS1 (ChIP) | A301-189A | Bethyl laboratories |
| NBS1 (ChIP-seq) | A300-289A | Bethyl laboratories |
| RNAPII (WB) | CTD4H8 | Millipore |
| RNAPII (ChIP, ChIP-seq) | F-12, sc-55492 | SCBT |
| RNAPII-ser5 (WB) | Ab5131 | Abcam |
| RNAPII-ser2 (WB and ChIP) | Ab5095 | Abcam |
| RNAPII-ser2 (ChIP-seq) | Ab5055 | Abcam |
| SNRNP200 | Sc-393170 | SCBT |
| PRP8 | Sc-55533 | SCBT |
| HNRNPM | Sc-20002 | SCBT |
| EFTUD2 | A300-957A | Bethyl laboratories |
| CCNT1 | A303-499A | Bethyl laboratories |
| CDK9 | A303-493A | Bethyl laboratories |
| TUBULIN | DM1A clone, T6199 | Sigma-Aldrich |
| Flag | M2 clone, F1804 | Sigma-Aldrich |
| HA | Sc-7392 | SCBT |
| Ku80 (ChIP) | Ab80592 | Abcam |
| IgG (mouse) | Sc-2025 | SCBT |
| IgG (rabbit) | Sc-2027 | SCBT |

**Table S4.** Sequences of deoxyoligonucleotides used in this study.

| **Primer** | **Forward (5’ to 3’)** | **Reverse (5’ to 3’)** |
| --- | --- | --- |
| GAPDH | CACATCGCTCAGACACCAT | GAGGTCAATGAAGGGGTCAT |
| RBM17 | TCCCACTTTCCCTTTCCCTC | CAAGGAGCACGAGGTTAAGC |
| ATXN7 | TACTTCGTCCTGACACCTGG | AAAGCATGAATGGGAAGCCG |
| Bambi | GGATTGACAGGAGCCCGG | CTCCGCGCCGTAAATAGC |
| HIV-1 LTR | GGGTCTCTCTGGTTAGA | GGGTTCCCTAGTTAGGC |
| ssODN MDC1 | GATGCTGTTTGTGTGAACCAATAATGAGGATAATTGATAATGTGTATCCTTCCCAGATCATGCAGGACTACAAGGACGACGATGACAAGCTCGATGGAGGATACCCCTACGACGTGCCCGACTACGCCGGAGGACTCGAGGAGGACACCCAGGCTATTGACTGGGATGTTGAAGAAGAGGAGGAGACAGAGCAATCCAGT | |
| PICh probe | ATGTCGACCTGCAGACATGGG (MS2.1) | |
| PICh probe | TGATCCTCATGTTTTCTAGGCAATTA (MS2.2) | |

**Table S5.** Double stranded shRNAs used in this study.

| **shRNA** | **Sequence (5’ to 3’)** |
| --- | --- |
| Con | cctaaggttaagtcgccctcgctcgagcgagggcgacttaaccttagg |
| MRE11 | ccggacgactgcgagtggactatagctcgagctatagtccactcgcagtcgttttttg |
| NBS1 | ccgggcaagcagatacatgggatttctcgagaaatcccatgtatctgcttgctttttg |
